# A computational approach to evaluate the combined effect of SARS-CoV-2 RBD mutations and ACE2 receptor genetic variants on infectivity: The COVID-19 host-pathogen nexus

**DOI:** 10.1101/2020.10.23.352344

**Authors:** Dana Ashoor, Noureddine Ben Khalaf, Maryam Marzouq, Hamdi Al Jarjanazi, Sadok Chelif, M. Dahmani Fathallah

## Abstract

SARS-CoV-2 infectivity is largely determined by the virus Spike protein binding to the ACE2 receptor. Meanwhile, marked infection rate differences were reported between populations and individuals. To understand the disease dynamic, we developed a computational approach to study the implications of both SARS-CoV-2 RBD mutations and ACE2 polymorphism on the stability of the virus-receptor complex. We used the 6LZG PDB RBD/ACE2 3D model, the mCSM platform, the LigPlot+ and PyMol software to analyze the data on SARS-CoV-2 mutations and ACE variants retrieved from GISAID and Ensembl/GnomAD repository. We observed that out of 351 RBD point mutations, 83% destabilizes the complex according to free energy (ΔΔG) differences. We also spotted variations in the patterns of polar and hydrophobic interactions between the mutations occurring in 15 out of 18 contact residues. Similarly, comparison of the effect on the complex stability of different ACE2 variants showed that the pattern of molecular interactions and the complex stability varies also according to ACE2 polymorphism. We infer that it is important to consider both ACE2 variants and circulating SARS-CoV-2 RBD mutations to assess the stability of the virus-receptor association and evaluate infectivity. This approach might offers a good molecular ground to mitigate the virus spreading.

## 1. Introduction

The emergent coronavirus, called Severe Acute Respiratory Syndrome Coronavirus 2 (SARS-CoV-2) has been spreading globally and efficiently since late 2019 (Elflein, 2020). SARS-CoV-2 is a new member of the coronaviruses family that includes SARS coronavirus (SARS-CoV) and Middle East Respiratory Syndrome coronavirus (MERS-CoV) (Chan et al., 2020). SARS-CoV-2 is the etiologic agent of the ongoing coronavirus disease (COVID-19) pandemic that claimed over two and a half millions life to date (EDCDP, 2020). The number of active cases per 100 thousand population estimates the prevalence of COVID-19 while the incidence is the number of new cases per time unit. The prevalence and incidence rate are good indicators for the tracking of infectious outbreaks (Noordzij et al., 2010). Indeed, they are very useful to evaluate the effectiveness of the mitigation efforts in the control of the disease spreading (Roser, 2020). Even though the spreading of the SARS-CoV-2 virus is global, the prevalence/incidence of the disease varies throughout the world, for some regions being more severely hit then others (EDCDP, 2020;Elflein, 2020;Roser, 2020). Even the overall case fatality ratio of COVID-19 varies between location and intensity of transmission (Lewnard and Lo, 2020;Ruan, 2020;Verity et al., 2020). Such discrepancies may be due to several factors that affect the virus infectious potency (infectivity) such as the health care prevention system organization and the stringency of the confinement measures (Koo et al., 2020), as well as the economic differences. Nevertheless, other important factors intrinsic to the virus structure and the host genetic background are crucial determinants of the SARS-CoV-2 infectivity. SARS-CoV-2 Invasion of human host cell is essentially but not exclusively mediated by the interaction of the virus capsid spike protruding S protein with the Angiotensin Convertase Enzyme 2 (ACE2) (Belouzard et al., 2012;Alifano et al., 2020). ACE2 is expressed on the surface of various human body cell-types (Walls et al., 2020;Zhao et al., 2020) where it displays a zinc metalloproteinase activity (Tipnis et al., 2000). The interaction ACE2/SARS-CoV-2 occurs through the receptor-binding domain (RBD) sequence of the S1 chain of the spike protein (Lan et al., 2020;Yan et al., 2020). This functional domain is highly prone to mutations (Nelson-Sathi et al., 2020). Meanwhile, the human ACE2 receptor gene that is located on chromosome X is highly polymorphic with hundreds of genetic variants identified to date in various populations (Cao et al., 2020). Noteworthy are the 1700 ACE2 variants reported in the Chinese populations though few variants are highly frequent and the majority of them are very rare (Lippi et al., 2020). Interestingly some ACE2 variants was shown to be significantly associated with the onset of diseases such arterial hypertension, diabetes mellitus, cerebral stroke, coronary artery disease, heart septal wall thickness and ventricular hypertrophy all considered as comorbidity factors of COVID-19 (Kowalczuk et al., 2008;Lu et al., 2012;Pinheiro et al., 2019;Calcagnile et al., 2020). The complexity of the SARS-CoV-2 S protein association with the ACE2 receptor is highlighted in a number of X-ray crystallography models. These are namely the 6LZG (SARS-CoV-2 Spike RBD /ACE2) complex, the 2AJF (SARS-CoV Spike RBD /ACE2) complex (Wang et al., 2020), the 6M0J (SARS-CoV-2 Spike RBD/ACE2) complex, and the 6VW1 (chimeric SARS-CoV/SARS-CoV-2 Spike RBD /ACE2) complex (Goodsell et al., 2020). These models have different resolution and are available at RCBS Protein Data Bank (https://www.rcsb.org/). Studies using these models have shown that the affinity of the RBD SARS-CoV-2 to the human ACE2 varies between ACE2 genetic variants suggesting that some people might be genetically protected where others are susceptible to COVID-19 (Lippi et al., 2020;Othman et al., 2020). Indeed, by measuring the ACE2 affinity for SARS-CoV-2 Spike using docking simulations, Calcagnile et al. (Calcagnile et al., 2020) showed that ACE2 variant S19P, which is more frequent in Africans and K26R more frequent in Europeans, are respectively protective and predisposing genetic factors to COVID-19. Meanwhile, other studies denied the association between ACE2 genetic polymorphism and the susceptibility to SARS infection (Chiu et al., 2004) and to COVID-19 (Lopera Maya et al., 2020). Nevertheless, the virus-host-cell interaction involves specific contacts between two structurally defined molecular entities. Therefore, it is very likely that the interaction between the SARS-CoV-2 and the human cell surface receptor ACE2 and thus the virus infectivity is determined by the genetic variations of both the pathogen and the receptor interacting sequences. This can reflect a higher magnitude of complexity in the interplay between this new viral pathogen and its human host.

In this paper, we report the development of a computational approach designed to examine this postulate. We first compiled the mutations observed in different isolates of the SARS-CoV-2 S protein RBD region as well as ACE2 receptor variants. Secondly, we assessed the stability of the complex between the virus RBD mutants and ACE2 isoform 1. Then we studied the structural basis of these molecular interactions. Comparison of the binding energy variations (ΔΔG) between the combinations of over 200 ACE2 genetic variants with 350 RBD mutated forms showed a gradient of stability among these combinations. We also observed significant variations in the patterns of polar and hydrophobic bonds between 18 RBD mutated contact residues and their partners at the interface of the complex with ACE2 Isoform 1.

We inferred from these data that both ACE2 receptor genetic polymorphism and the circulating SARS-CoV-2 RBD genetic type determine the stability of the virus-receptor association and might affect the infectivity of SARS-CoV-2. Furthermore, these molecular interaction patterns highlight the level of complexity of the interplay between SARS-CoV-2 and its major ACE2 receptor. The data also point out to the benefit this computational approach may bring to the rapid evaluation of the infectivity of a given viral genetic subtype in consideration of an individual or a group dominant ACE 2 variant. This might help fine-tuning the prevention strategies against the spreading of the virus.

## 2. Materials and Methods

### 2.1. SARS-CoV-2 S protein RBD mutations and ACE2 genetic variants data retrieval

We first retrieved the mutations located at the SARS-CoV-2 Spike protein RBD region (351 missense mutations), Wuhan strain (NCBI Reference: YP_009724390.1). The data was obtained from the SARS-CoV-2 Spike protein mutation table compiled by the Singapore Bioinformatics Institute (SBI) at (https://mendel.bii.a-star.edu.sg/METHODS/corona/beta/MUTATIONS/hCoV-19_Human_2019_WuhanWIV04/hCoV-19_Spike_new_mutations_table.html) based on sequence information provided by GISAID initiative (https://www.gisaid.org/). Data for genetic variants of ACE2 was from the Ensembl dbSNP, GnomAd and UNIPROT databases.

To identify the amino acids differences in the spike RBD between the new Wuhan strain SARS-CoV-2 and previously known SARS-CoV (NCBI Reference: YP_009825051.1), the sequence of the RBD domain of both viruses was determined using Pfam sequence analysis (https://pfam.xfam.org) of the S full -protein. SARS-CoV-2 RBD protein sequence was aligned with SARS-CoV RBD protein using Clustal Omega alignment tool (https://www.ebi.ac.uk/Tools/msa/clustalo/). The locations of the variant amino acids were identified and the reported mutations of SARS-CoV-2 at these locations were examined for possible reversed mutations to the SARS-CoV sequence. The effect of these mutations on the complex stability with the human ACE2 receptor was examined as described in the next section.

### 2.2. RBD mutation effect analysis

The PDB structure (PDB ID 6LZG) of the complex formed between the SARS-CoV-2 Spike protein RBD and the human ACE2 receptor isoform 1 was downloaded from the RCSB database (https://www.rcsb.org). ACE2 isoform 1 sequence correspond to the ACE2 sequence that was used to generate the 6LZG 3D model. This structural complex model was taken as reference model for this study. The structure was cleaned from water and heteroatoms using PyMol software (PyMol Molecular). The mCSM server, a web server (http://biosig.unimelb.edu.au/mcsm/) that predicts the effects of mutations in proteins using graph-based signatures platform (Pires et al., 2020) was used to evaluate the effect of the mutations reported in the RBD domain on the stability of the virus-receptor complex. The prediction relies on graph-based signatures that encode distance patterns between atoms and are used to represent the protein residue environment and to train predictive models. The models predict the stability changes through free energy (ΔΔG) difference due to a single nonsynonymous substitution in the amino acid chain of the protein-protein complex. Finally, mutations were ranked by their ΔΔG values. Site Directed Mutator server (SDM) (http://marid.bioc.cam.ac.uk/sdm2/) (Worth et al., 2011) was used to create the 23 contact residues mutants and chain replacement was used to build the complexes using PyMol (Schrödinger, 2010;Verity et al., 2020). Interacting residues in the generated structures, involving polar and hydrophobic interactions from both receptor and ligand, were determined using LigPlot+ software (Laskowski and Swindells, 2011) and PyMol software was used to illustrate the interacting interface in presence of the most destabilizing mutations in the RBD.

### 2.3. hACE2 variants analysis

The protein sequence for human ACE2 Isoform 1 (hACE2) was downloaded from the Uniprot data bank (https://www.uniprot.org/) (Uniprot ID: Q9BYF1-1) and used as a reference sequence in this study. We analyzed 231 human ACE2 SNPs causing protein sequence variations collected from the Ensembl dbSNP,GnomAD and UNIPROT databases. We used the PDB structure (PDB ID 6LZG), LigPlot+ software (Laskowski and Swindells, 2011), and mCSM server (http://biosig.unimelb.edu.au/mcsm/) to identify possible amino acid substitutions that affect the virus-receptor complex stability, and their interaction pattern. We then built protein complexes (hACE2/RBD) using SDM server and Pymol software to analyze S protein mutations effect on complex stability for selected hACE2 variants with predicted destabilizing effect on the original complex. We used LigPlot+ and PyMol softwares to analyze the molecular interactions at the complex interface.

## 3. Results

### 3.1. SARS-CoV-2 S protein RBD mutations retrieval

This study focused on the investigation of the structural variations of the Spike’s RBD of the SARS-CoV-2 virus from the first reported Wuhan strain (NCBI Reference: YP_009724390.1). Alignment of the SARS-CoV surface glycoprotein and SARS-CoV-2 spike RBD sequence showed 74% identity and one insertion (V483) (data not shown). Random search of the GISAID data bank yielded 351 non-synonymous mutations with some positions changing from one up to eight different residues as shown in Figure 1. Out of these mutations, 24 are reverse mutations to the SARS-CoV RBD sequence.

**Figure 1.**
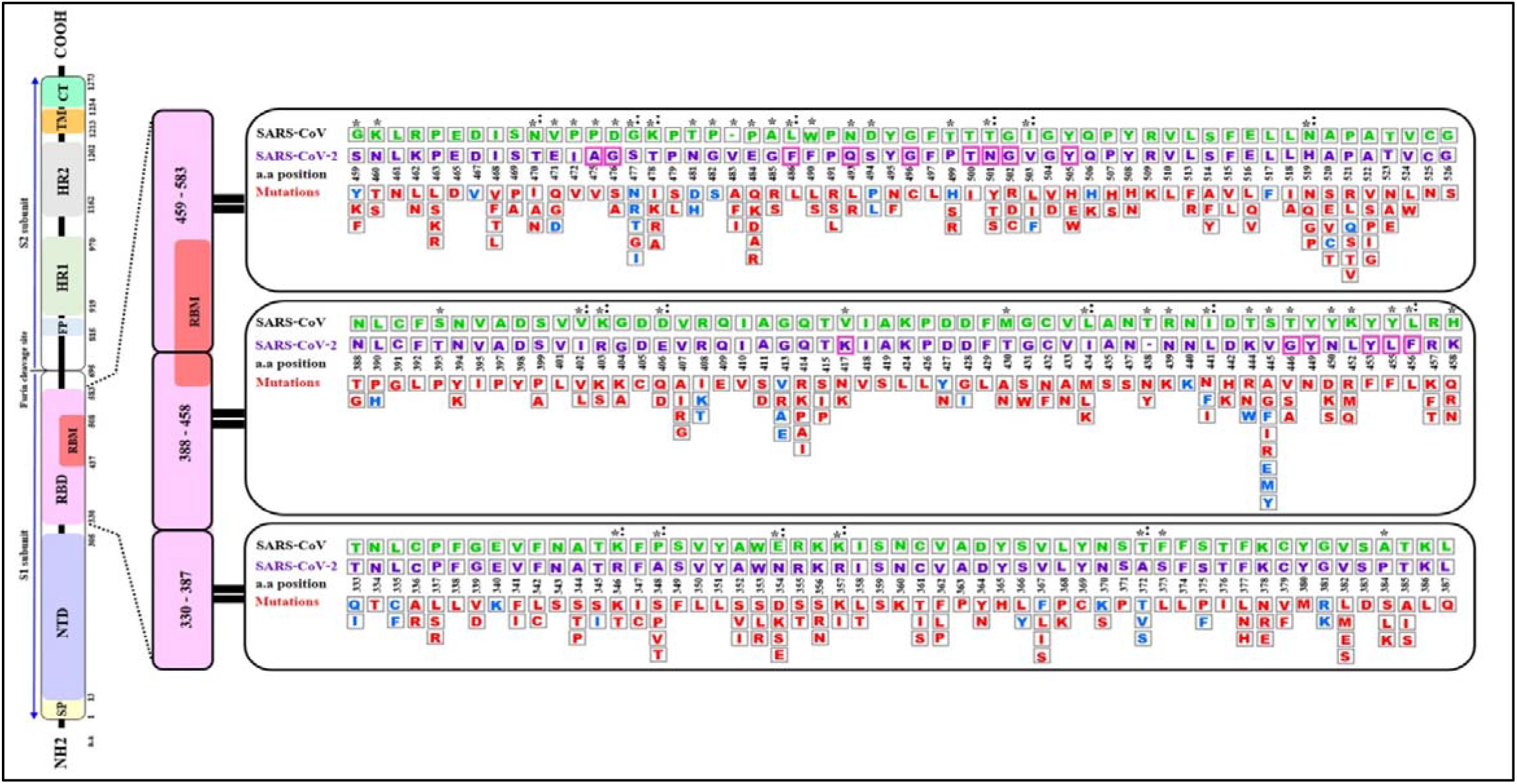
A Schematic diagram of the of SARS-CoV-2 S spike protein sequence (purple letters) and the mutations in the Receptor Binding Domain (RBD) aligned with SARS-CoV sequence (green letters). The S protein is cleaved into S1 and S2 subunit at the polybasic furin cleavage site. Stabilizing and destabilizing mutations of the RBD domains are represented with blue and red letters respectively. Asterisks display amino acid variations between SARS-CoV-2 and SARS-CoV RBDs. Magenta boxes highlighted the positions of the 15 (out of 18) contact residues where mutations were reported. Colons indicate reverse mutations to SARS-CoV sequence. Empty box with dashes (-) represent an insertion of a valine (V) at position 483 of SARS-CoV-2. Signal peptide (SP), N-terminal domain (NTD), Receptor-Binding Domain (RBD), Receptor-Binding Motif (RBM), Fusion Peptide (FP), Heptad Repeat 1 (HR1), Heptad Repeat 2 (HR2), Transmembrane Domain (TM) and Cytoplasmic Tail (CT) a.a (amino acid). The RBD sequence analyzed in this diagram corresponds to residues 333-527 of the S protein RBD in the structure PDB ID 6LZG.

### 3.2. Analysis of the RBD mutations effect on the S protein/ACE2 complex stability

The PDB ID 6LZG model of the complex RBD S protein (Wuhan strain sequence)/Human ACE2 isoform 1 was selected for this study since it displayed the best resolution (2.5 Å) of the complexes available in the PDB database. The LigPlot+ analysis of the interactions showed that there are 18 residues of SARS-CoV-2 interacting with the hACE2 receptor, (Supplementary Table 1), with ten of them forming polar bonds with the receptor. In addition, these 18 contact residues have one or more hydrophobic interactions with the hACE2 receptor except for G446 as it forms only one polar interaction with Q42 of the ACE2 receptor. A total of 12 polar bonds and 65 hydrophobic interactions were identified at the complex interface as shown in Figure 2. These interactions involve the 18 contact residues (Supplementary Table 1). Out of these residues, 15 showed multiple mutations as highlighted in Figure 1.

**Figure 2.**
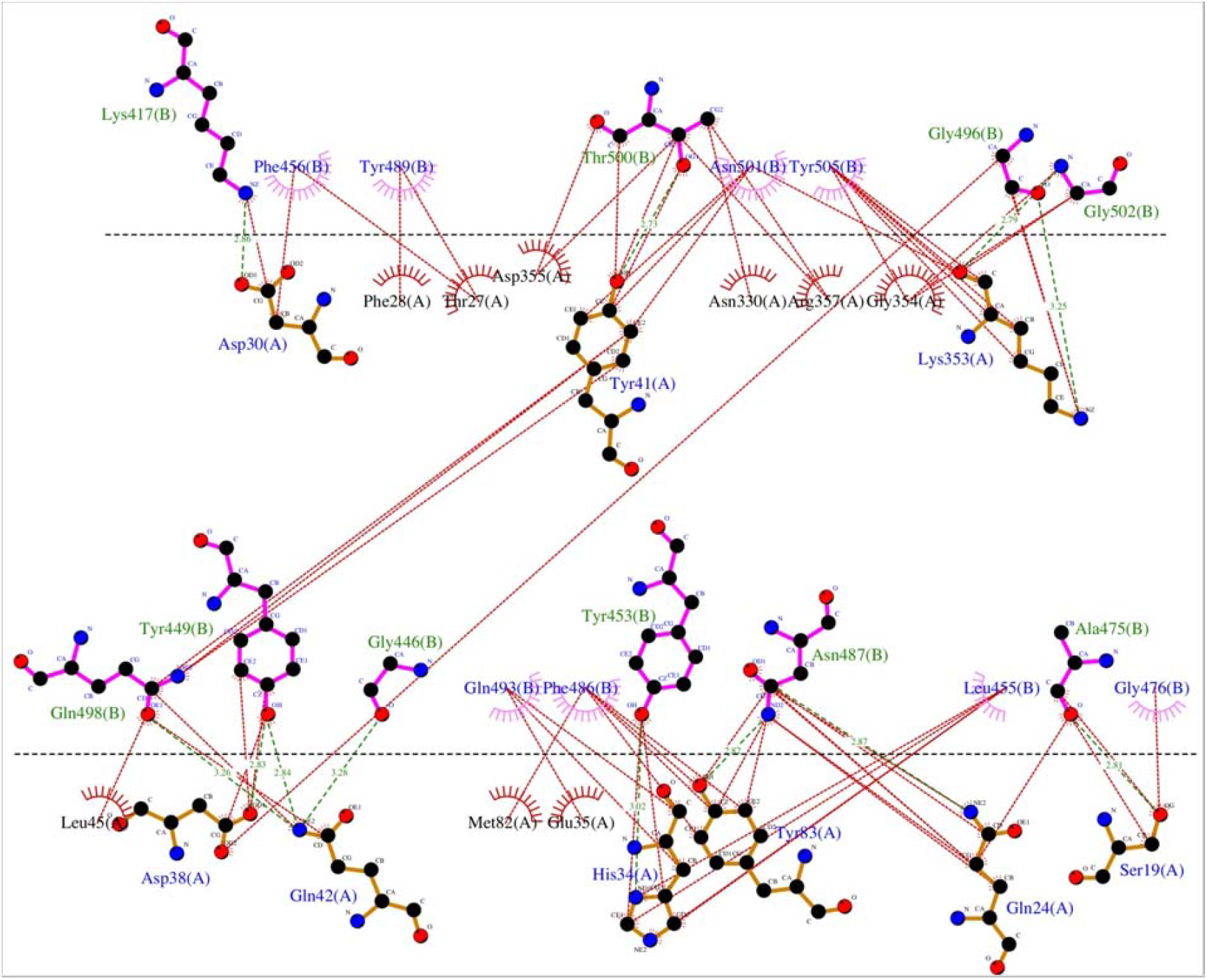
Representation of the polar and hydrophobic interactions between SARS-CoV-2 spike RBD of the Wuhan strain (chain B) and hACE2 isoform 1 receptor (chain A) of the complex structure (PDB ID 6LZG). The dotted green lines show the polar interactions, and the dotted red lines the hydrophobic interactions.

The complete set of data on the effect of the SARS-CoV-2 RBD 351 mutations on the complex stability are displayed in Supplementary Material 1. Figure 3 shows the distribution of the Free energy difference of all analyzed mutations. We observe that, out of the 351 mutations reported in the GISAID server, 83% of the mutations were classified as “Destabilizing”, and 17% were classified as “Stabilizing” according to their ΔΔG value. Energy difference values ranged from 0.788 Kcal/mol (T333Q, most stabilizing) to −2.482 Kcal/mol (G502R, most destabilizing). Among the destabilizing mutations, three mutations were classified as “Highly destabilizing” with ΔΔG values less than −2.0 Kcal/mol (G502R, N501Y and Y505E) (Supplementary Material 1). Furthermore, this analysis showed that among the mutations that reversed to SARS-CoV RBD sequence, mutation A372T is predicted to have a relatively good stabilizing effect on the complex with hACE2 isoform 1 as shown by the value of the ΔΔG (+0.508) Figure 2

**Figure 3.**
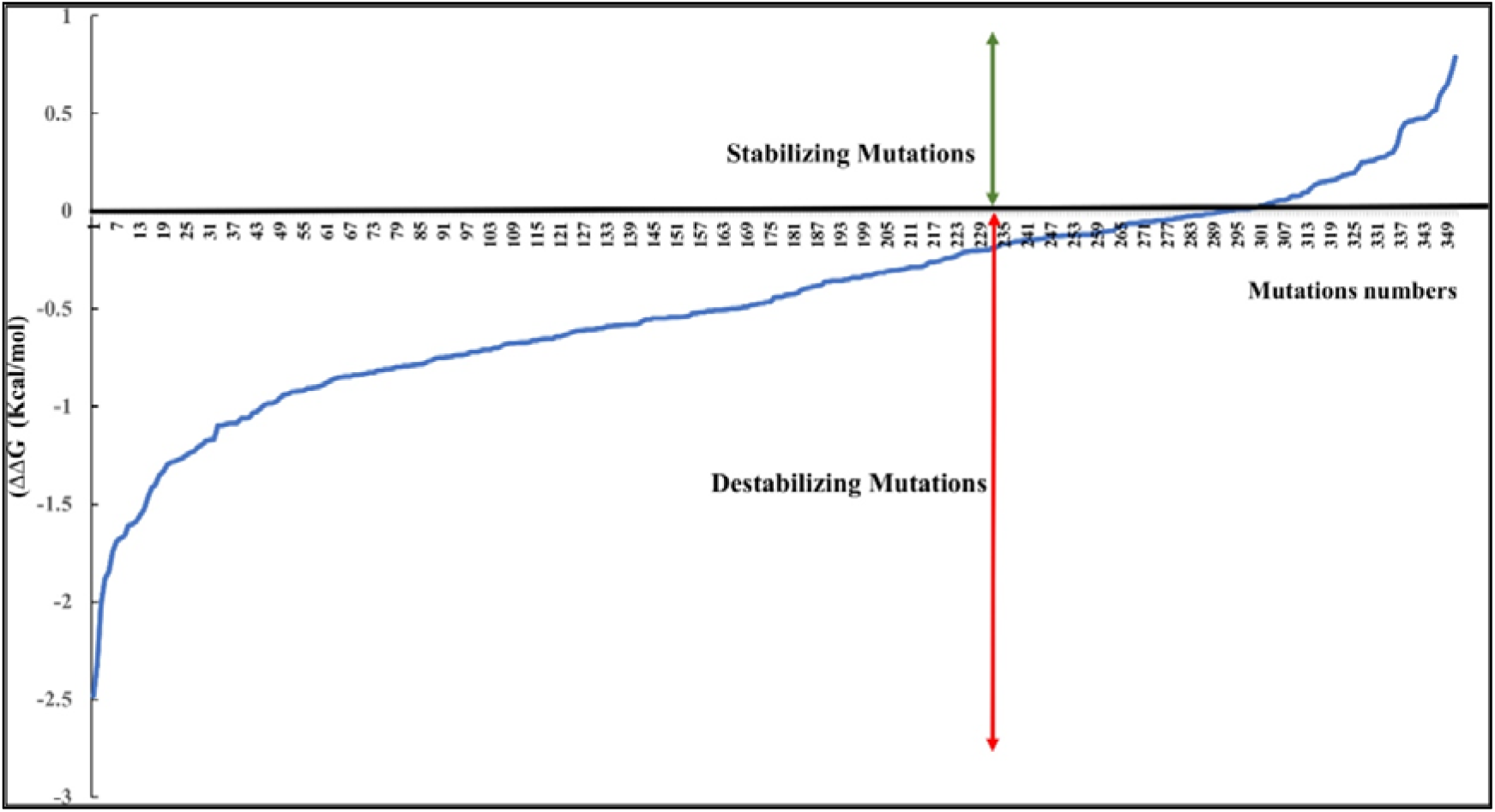
Distribution of the free energy ΔΔG differences of the reported 351 nonsynonymous substitution in the protein sequence of the hACE2-SARS-COV-2 S RBD complex.

SDM generated complexes for the S protein mutants were used to establish the molecular interaction patterns using LigPlot+. Out of the mutations affecting the 18 interacting residues only six mutations (K417N, G446A, Y449N, Y453R, Q493R, and T500I), predicted as destabilizing mutations, cause polar bond differences (Supplementary Table 1). Three out of these residues were shown to be conserved residues between SARS-CoV-2 and SARS-CoV (Y449, Y453, and T500). Figure 4 shows the differences in polar interactions between theses mutants and the human ACE2 receptor as compared to the reference complex. Inter-chain polar bonds are shown to be missing in all the mutated structures at the complex interface level except for Q493R, as a new polar interaction with E35 of ACE2 receptor was introduced. For the hydrophobic interactions, the most stabilizing and destabilizing mutations were analyzed (40 mutants) and the results showed that the interaction patterns did not change for the mutations that occurred out of the contact residues. Figure 5 shows the hydrophobic interaction changes in comparison to Wuhan SARS-CoV-2 strain for the Spike/ACE2 complex for mutations that occurred in contact residues. Mutations in contact residue N501, G502 and Y505 were shown to be the most destabilizing mutations according to ΔΔG predictions. Ranked from the highest destabilizing to the lowest these mutations are G502R, N501Y, Y505E, N501T and N501S with the following respective ΔΔGs (−2.482, −2.317, −2.014, −1.689, −1.589 Kcal/mol). The least destabilizing mutant among the contact residues is L455F with a ΔΔG of −0.124 Kcal/mol. Interestingly, all of these mutants have no missing polar interactions. However, they contain extra hydrophobic interactions compared to the original complex. Polar bonds seem to be important for maintaining the complex stability between RBD and hACE2 receptor, which explain the high-energy difference, predicted by mCSM, however gaining extra hydrophobic interactions highly affect the complex stability (Supplementary Table 1).

**Figure 4.**
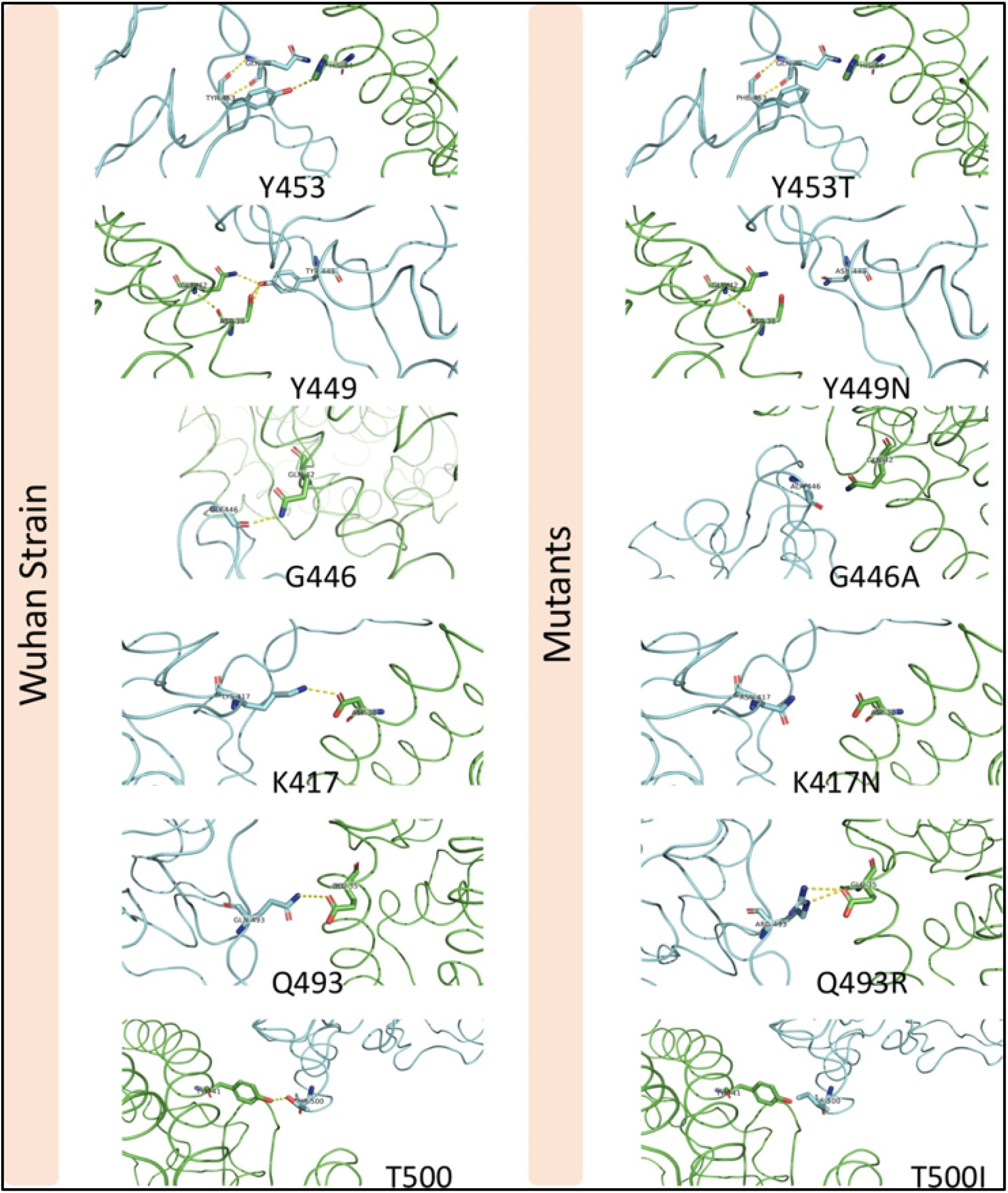
Mutations in the SARS-COV-2 RBD sequence causing changes in the pattern of polar interactions with the human ACE2 receptor. Original and mutated residues (K417N, G446A, Y449N, Y453R, Q493R, and T500I) are represented in sticks. Protein chains are shown in cartoon representation (RBD; Blue / hACE2; Green). Polar interactions are shown with yellow dashes.

**Figure 5.**
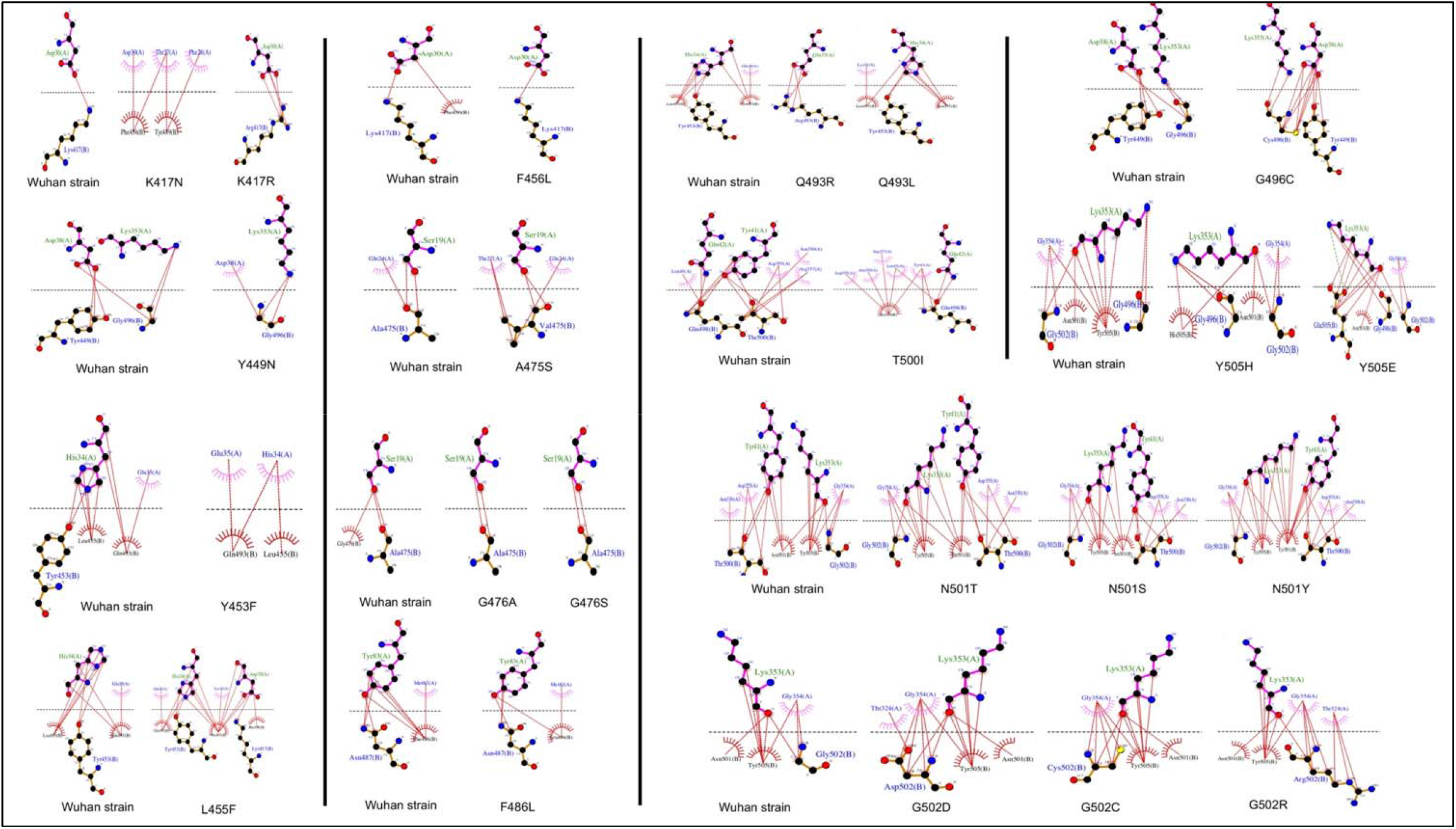
Representation of the hydrophobic interaction of SARS-CoV-2 spike RBD (chain B) with hACE2 receptor (chain A) of the 6LZG complex. Hydrophobic bonds are represented in Red lines and Salt bridges in dotted green lines.

### 3.3. Analysis of the effect of ACE2 genetic polymorphism on the S protein/ACE2 complex stability

The analysis of the effect of hACE2 variants on the SARS-CoV2/hACE2 complex stability showed that out of 231 variants, 203 (88%) are destabilizing (Supplementary Material 2). Out of these, six variants (D355A, W163R, D355N, G405E, G377E and Y252N) have highly destabilizing effect with a ΔΔG less than −2.0 Kcal/mol (Supplementary Material 2).

The analysis of the hACE2 residues involved in the interaction of the receptor with the SARS-CoV-2 Spike’s protein, showed that variants D355A, D355N, E35K, F40L, E35D, M82I, S19P and T27A are destabilizing variants (Supplementary Material 2), with the D355A and D355N, predicted as being highly destabilizing (−2.775 Kcal/mol and-2.342 Kcal/mol, respectively). Interaction analysis showed that D355 establishes a network of intramolecular polar interactions with hACE2’ Y41, G326, N331, L351 and R357 in addition to the polar interaction with the Spike’ T500 Figure 6 A. Meanwhile, variant D355A Figure 6 B causes the loss of this interaction network, suggesting that this mutation strongly destabilizes the complex, which is in line with the ΔΔG prediction. The D355N mutation interaction analysis Figure 6 C shows that N355 maintains only three polar bonds with hACE2 R357, Y41 and Spike T500.

**Figure 6.**
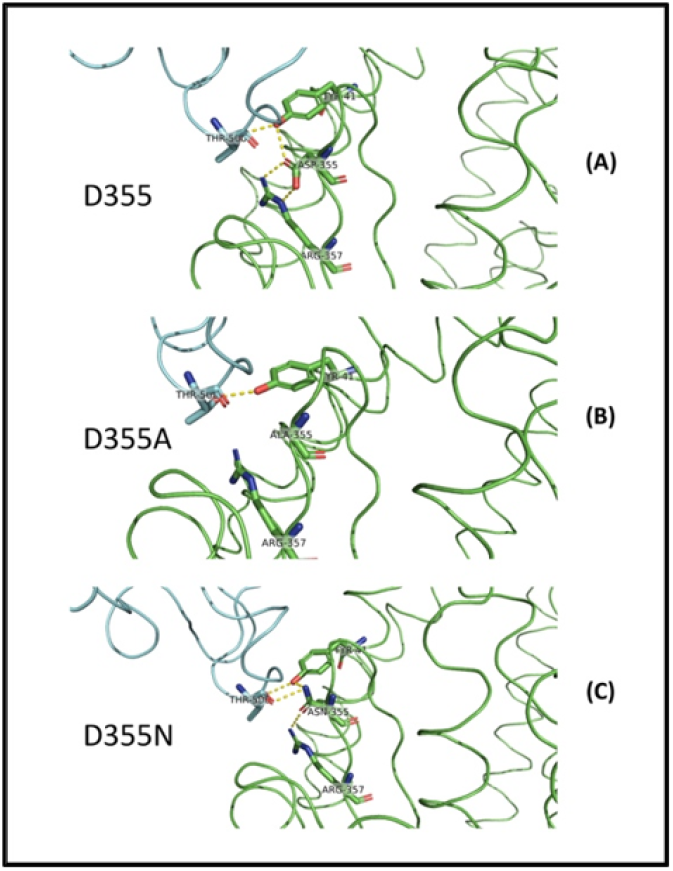
Genetic variations in the human ACE2 receptor sequence causing changes in the pattern of polar interactions with the SARS-COV-2 Wuhan strain RBD sequence. Original D355 (A) and variant D355A (B) and D355N (C) are shown in sticks representation. Protein chains are shown in cartoon representation (RBD; Blue / hACE2; Green). Polar interactions are represented by yellow dashes.

The eight hACE2 variants in residues involved in the interaction interface of the receptor with the SARS-CoV-2 Spike’s protein were screened for their effect against all the reported S protein mutations (Supplementary Material 2). Results showed a significant change in energy prediction (threshold set at 20% change in ΔΔG) for some variants against Spike mutants compared to the hACE2 isoform 1, as shown in Figure 7. In fact, D355N was predicted to have more destabilizing effect with mutants N501T, N501S and G502D with ΔΔG values reduced by 34, 39 and 22% respectively, compared to hACE2 isoform 1. In addition, the variant is predicted to be less destabilizing with the mutant G502C with a ΔΔG value increased by 37%. Similarly, variants T27A, M82I, and S19P show a less destabilizing effect with Spike mutants F456L, F486L and G476S/A, respectively.

**Figure 7.**
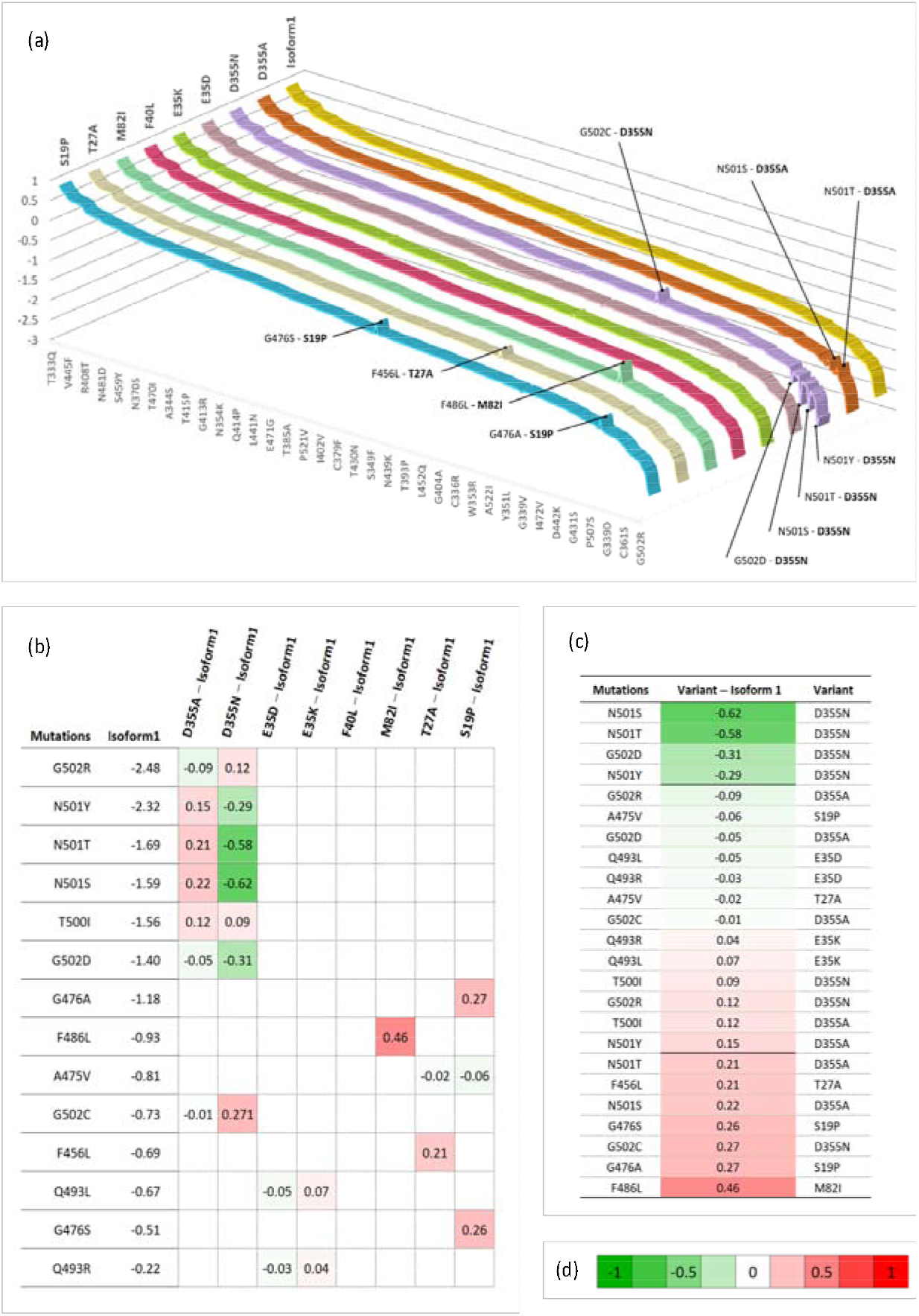
Gradient of binding stability between SARS-CoV-2 mutants and ACE2 variants at the contact residues. **(a)** 3D ribbon chart showing the ΔΔG distribution depending on mutations and variants. Each ribbon represents the values of ΔΔG for a specific variant (bold labels) with the different mutants. For a given line, most values are equal to Isoform1 variant, which drives the trend (yellow ribbon). The chart shows the spikes having a distance ≥ 0.2 to Isoform1. The value of 0.2 is selected empirically for visual convenience and as a threshold to label a spike (combination mutation/ACE2 variant) as important. **(b)** Matrix showing the ΔΔG difference between Variants and Isoform1 for various contact residues mutations. The difference represents an algebraic measure ranging from −0.62 to 0.46. The matrix cells are colored using color gradient (d) to illustrate the intensity and direction of the gap. The matrix lines represent the mutations with at least one non-null difference. The non-null values are shown inside the cells and can be used to assess the difference with Isoform1. **(c)** Table showing the difference between variants ΔΔG and Isoform1 ΔΔG for specific mutations in term of a unidimensional scale. It is obtained by discarding all null cells and by sorting remaining cells in ascending order. A bold line is used to distinguish important (having a distance ≥ 0.2 to Isoform1). **(d)** A gradient palette to visually weight a continuous change from −1 (Green) to +1(Red) through 0 (White). The choice of numerical limits (−1 and +1) is made to facilitate the visual reading of the intensity and sense of the gap to Isoform1.

Table 1 summarizes the variations of molecular interactions numbers and patterns between the SARS-CoV-2 RBD mutations occurring in the 18 residues the virus engages to interact with the ACE2 isoform1 and the eight variants found to be significantly destabilizing when tested against the Wuhan RBD sequence in comparison to hACE2 isoform1. The number and position of polar and hydrophobic bonds in addition to the salt bridges characterize each interaction pattern. The most striking interactions variations concern SARS-CoV-2 RBD mutants N501T and N501S and ACE2 variant D355N. Interactions with corresponding residues are detailed in Supplementary Table 2.

**Table 1.**
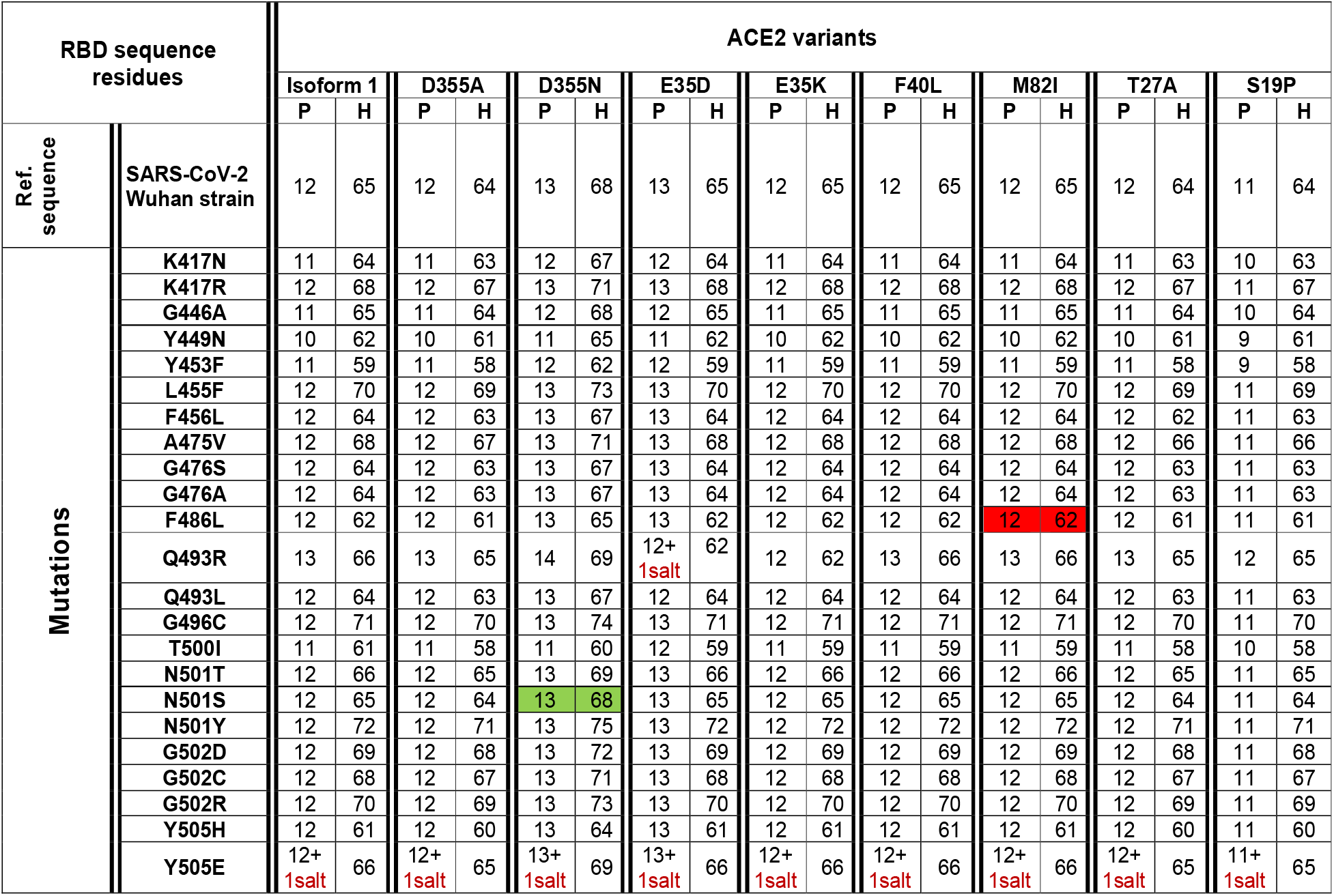
Variation of the number of polar and hydrophobic interactions between 24 different SARS-CoV-2 spike RBD mutations that occurred in the 18 contact residues with nine hACE2 genetic variants. P: polar, H: hydrophobic. The red and the green show respectively the most and the least divergent interaction from the Wuhan virus RBD sequence and ACE2 isoform1 interaction considering both the ΔΔG and the pattern of molecular interactions.

## 4. Discussion

As the COVID-19 pandemic progresses, it is becoming more and more important to understand the molecular basis of viral infectivity of the SARS-CoV-2, the etiologic agent of the disease. This is crucial to evaluate correctly the virus capacity to invade the host cell and exploit its molecular machinery to replicate and produce infectious viral progeny that may invade other cells and lead to disease progression (Yan et al., 2020). It is also important to investigate the difference in susceptibility to the infection considering the differences observed in the disease prevalence between various regions of the world as well as inter individual variability (Hoffmann et al., 2020;Hou et al., 2020;Hussain et al., 2020). This necessitates a multidisciplinary approach that would be the most appropriate to get the best insights into SARS-CoV-2 infectivity. These insights will likely guide the appropriate response of public health authorities worldwide to combat the pandemic (Frieden and Lee, 2020;Hellewell et al., 2020). The genetic variability of a specific virus receptor that affect its infective power was demonstrated for the human immunodeficiency virus (HIV). Indeed, in humans the CD4 receptor variant (C868T) and the CCR5 receptor CCR5-Δ32 variant confer respectively susceptibility and resistance against certain HIV strains (Marmor et al., 2006). Other studies on group 2 coronavirus that causes hepatitis in mouse showed that the viral receptor allelic variants were associated with altered virus binding and hence with host susceptibility (Ohtsuka and Taguchi, 1997). Similarly, it was reported that rs73635825 (S19P) and rs143936283 (E329G) alleles of the hACE2 receptor may offer some level of resistance/protection against the infection because of a low affinity to the virus RBD region (Calcagnile et al., 2020;Hussain et al., 2020). Nevertheless, most of the accumulated evidences suggesting that genetic polymorphism of the ACE2 gene may be associated to susceptibility or protection from SARS-CoV-2 (Chiu et al., 2004;Calcagnile et al., 2020;Lippi et al., 2020) are based on comparative studies of the molecular interactions between ACE2 variants and a single genetic type of the virus. Hence, not considering the high number of mutations in the RBD region of the virus. In this work, we have considered both the SARS-CoV-2 variability introduced by spontaneous mutagenesis and the natural genetic variation of the ACE2 receptor (Li et al., 2020). Indeed, we have compiled 351 non-synonymous natural mutations that occurred in the SARS-CoV-2 Wuhan strain S protein RBD region. This high number of mutations in a relatively short time suggest a genetic instability of the virus receptor-binding domain knowing that RNA-viruses have generally low mutagenesis rate. We showed that over eighty percent of these mutations destabilizes the complex with ACE2 isoform 1. This data suggests that the crushing number of SARS-CoV-2 naturally occurring genetic changes are most likely making the virus less infective. We also found that 24 mutations are reverse mutations to the SARS-CoV RBD sequence. Fourteen of these reversed mutations showed a destabilizing effect on the SARS-CoV-2 RBD/hACE2 complex while four of them were stabilizing mutations. This would partially explain the difference in infectivity of SARS-CoV and SARS-CoV-2 observed from epidemiological and clinical data (EDCDP, 2020).

On another hand, studies of SARS-CoV-2 RBD/ACE2-isoform 1 complexes focusing on the mutations in the RBD contact residues showed that only six mutants (K417N, G446, Y449N, Y453R, Q493 and T500I), predicted as destabilizing mutations, cause polar bond changes. Previous studies reported 17 residues in the SARS-CoV-2 RBD to be involved in the interaction with the ACE2 receptor. Fourteen of these residues are conserved between SARS-CoV and SARS-CoV-2 with eight residues being contact residues (Lan et al., 2020;Walls et al., 2020). In this study, we found that 18 residues in the RBD region are involved in the interaction with the ACE2 receptor. The interaction pattern did not change for the mutations that occurred in non-contact residues, which suggest that these residues act as a structural scaffold that is important for the virus binding to its receptor. Polar bonds seem to be important for maintaining the complex stability between RBD and hACE2 receptor, which explain the high-energy difference, predicted by mCSM. In addition, out of the 18 described contact residues, ten different mutations occurring in conserved residues between SARS-CoV-2 and SARS-CoV (Y449N, Y453R, G496C, T500I, G502D, G502C, G502R, Y505H, Y505E, and Y505W) are predicted to be destabilizing mutations. This is consistent with the fact that conserved residues are important to maintain the overall correct RBD domain three-dimensional fold needed for efficient binding to the ACE2 receptor. The loss of polar bonds constitutes an important factor in protein-protein interaction.

Besides, the analysis of the effect of hACE2 variants on the SARS-CoV-2 Wuhan Strain/ACE2 complex stability showed that out of 231 variants, 88% are predicted to be destabilizing. Eight ACE2 variants of ACE2 residues involved in the interaction interface of the receptor with the SARS-CoV-2 Spike’s protein are destabilizing. Variants D355A and D355N predicted as highly destabilizing are potentially protective which is consistent with previously reported experimental data. This in silico study of the molecular bonds showed that D355 establishes a network of polar interactions with ACE2 residues Y41, G326, N331, L351 and R357 and Spike’ T500. The substitution of the 355 aspartic acid by an alanine (D355A) causes the loss of this interaction network. 292, mutation D355N shows that Asparagine 355 maintains only three polar bonds with hACE2 R357, Y41 and Spike T500, which likely makes this variant the most, destabilizing one of the RBD/ACE2 complex of the RBD mutations studied. This result suggest that this variant is potentially more resistant to the virus attachment. In addition, in our study ACE2 variant S19P more frequent in Africans and reported to be a protective genetic variant (Calcagnile et al., 2020), showed the lowest number of interactions with the different spike RBD mutants Table 1. Therefore, if we consider our findings together with the literature report on the genetic susceptibility to SARS-CoV-2 infection, we can argue that there is no absolute susceptible or protective ACE2 genetic variant. Indeed, the same given ACE2 variant might protect from a virus genetic type and be susceptible to another depending on the molecular interactions engaged between the genetic type of the circulating virus and an individual’s receptor genetic variant. This may explain the striking individual difference observed globally in clinical presentations with some patients remaining asymptomatic, whilst others developing the disease (COVID-19) considering that all have the same comparable baseline risk. Hence, using a computational approach to assess SARS-CoV-2 infectivity through the evaluation of the binding of the circulating genotype with individual or population-dominant ACE2 variant could be a rapid and reasonable way to estimate the spreading of the disease. Indeed, experimental measurement of the S protein-ACE2 affinity is time consuming and cannot be deployed for a large number of RBD mutants-ACE2 variants combinations. Nevertheless, it should be noted that accumulation of several mutations in the same virus type have usually cumulative effects that cannot be measured by the approach we used in this work unless a more refined RBD/ACE structure with high resolution [less than 2 Angstrom] becomes available. Therefore, the approach developed in this work can be applied using a platform software such as MutaBind2 (https://lilab.jysw.suda.edu.cn/research/mutabind2/). Furthermore, combining in silico information with clinical and other conventional epidemiological data could be very helpful in stratifying the risk of infection and for fine-tuning of the mitigation efforts to circumvent disease progress.

## Author Contributions

Dana Ashoor and Noureddine Ben Khalaf: In Silico analysis, methodology, data curation, writing and editing (both authors contributed equally to this work). Maryam Marzouq: Mutations’ review, Illustrations figures and tables, revision. Hamdi Al Jarjanazi: primary and secondary data retrieval formatting and management. Sadok Chlif: Illustrations and mathematical data analysis. M. Dahmani Fathallah: Project conception, work design, data analysis, writing, editing, and supervision. All authors have read and agreed to the submitted version of the manuscript.

