## Supplementary Material 3 for "A computational approach to evaluate the combined effect of SARS-CoV-2 RBD mutations and ACE2 receptor genetic variants on infectivity: The COVID-19 host-pathogen nexus"

**Supplementary Table 1**: Changes in the polar (red P) and hydrophobic (green H) interactions between spike RBD and hACE2 receptor contact resides in different mutated strains of SARS-CoV-2. Note: only missing, changed and new interactions are mentioned under each mutation, all other interactions are same as the Wuhan strain.10 out of 18 contact residues of spike RBD show polar interactions with hACE2 receptor (highlighted in red). (*)= salt bridge

| Spike RBD Wuhan strain contact residues | hACE2 contact residues | Mutants | | | | | | | | | | | | |
|---|---|---|---|---|---|---|---|---|---|---|---|---|---|---|
| | | K417N | K417R | G446A | Y449N | Y453F | L455F | F456L | A475V | G476S | G476A | F486L | Q493R | Q493L |
| Lys417 | 1X Asp30(P)<br>1X Asp30(H) | Missing<br>Missing | 4X Asp30(H) | | | | | | | | | | | |
| Gly446 | 1X Gln42(P) | | | Missing | | | | | | | | | | |
| Tyr449 | 1X Asp38(P)<br>1X Gln42(P)<br>3X Asp38(H) | | | | Missing<br>Missing<br>Missing | | | | | | | | | |
| Tyr453 | 1X His34(P)<br>2X His34(H) | | | | | Missing<br>Missing | | | | | | | | |
| Leu455 | 4X His34(H) | | | | | 1X His34(H) | 4X Asp30(H)<br>1X Lys31(H) | | | | | | | |
| Phe456 | 1X Thr27(H)<br>1X Asp30(H) | | | | | | | Missing | | | | | | |
| Ala475 | 1X Ser19(P)<br>2X Ser19(H)<br>1X Gln24(H) | | | | | | | | 2X Gln24(H)<br>2X Thr27(H) | | | | | |
| Gly476 | 1X Ser19(H) | | | | | | | | | Missing | Missing | | | |
| Phe486 | 1X Met82(H)<br>4X Tyr83(H) | | | | | | | | | | | 1X Tyr83(H) | | |
| Asn487 | 1X Gln24(P)<br>1X Tyr83(P)<br>6X Gln24(H)<br>3X Tyr83(H) | | | | | | | | | | | | | |
| Tyr489 | 1X Thr27(H)<br>1X Phe28(H) | | | | | | | | | | | | | |
| Gln493 | 2X His34(H)<br>1X Glu35(H) | | | | | 1X His34(H) | | | | | | | 1X Glu35 (P)<br>Missing<br>4X Glu35(H) | 1X His34(H)<br>Missing<br>1X Lys31(H) |
| Gly496 | 1X Lys353(P)<br>1X Asp38(H)<br>2X Lys353(H) | | | | | | | | | | | | | |
| Gln498 | 1X Gln42(P)<br>3X Tyr41(H)<br>2X Gln42(H)<br>1X Leu45(H) | | | | | | | | | | | | | |
| Thr500 | 1X Tyr41(P)<br>3X Tyr41(H)<br>1X Asn330(H)<br>2X Asp355(H)<br>2X Arg357(H) | | | | | | | | | | | | | |
| Asn501 | 3X Tyr41(H)<br>1X Lys353(H) | | | | | | | | | | | | | |
| Gly502 | 1X Lys353(P)<br>1X Lys353(H)<br>2X Gly354(H) | | | | | | | | | | | | | |
| Tyr505 | 5X Lys353(H)<br>1X Gly354(H) | | | | | | | | | | | | | |

**Supplementary Table 1**: (Continued)

| Spike RBD Wuhan strain contact residues | hACE2 contact residues | Mutants | | | | | | | | | |
|---|---|---|---|---|---|---|---|---|---|---|---|
| | | G496C | T500I | N501T | N501S | N501Y | G502D | G502C | G502R | Y505H | Y505E |
| Lys417 | 1X Asp30(P)<br>1X Asp30(H) | | | | | | | | | | |
| Gly446 | 1X Gln42(P) | | | | | | | | | | |
| Tyr449 | 1X Asp38(P)<br>1X Gln42(P)<br>3X Asp38(H) | | | | | | | | | | |
| Tyr453 | 1X His34(P)<br>2X His34(H) | | | | | | | | | | |
| Leu455 | 4X His34(H) | | | | | | | | | | |
| Phe456 | 1X Thr27(H)<br>1X Asp30(H) | | | | | | | | | | |
| Ala475 | 1X Ser19(P)<br>2X Ser19(H)<br>1X Gln24(H) | | | | | | | | | | |
| Gly476 | 1X Ser19(H) | | | | | | | | | | |
| Phe486 | 1X Met82(H)<br>4X Tyr83(H) | | | | | | | | | | |
| Asn487 | 1X Gln24(P)<br>1X Tyr83(P)<br>6X Gln24(H)<br>3X Tyr83(H) | | | | | | | | | | |
| Tyr489 | 1X Thr27(H)<br>1X Phe28(H) | | | | | | | | | | |
| Gln493 | 2X His34(H)<br>1X Glu35(H) | | | | | | | | | | |
| Gly496 | 1X Lys353(P)<br>1X Asp38(H)<br>2X Lys353(H) | 7X Asp38(H) | | | | | | | | | |
| Gln498 | 1X Gln42(P)<br>3X Tyr41(H)<br>2X Gln42(H)<br>1X Leu45(H) | | 2X Tyr41(H) | | | | | | | | |
| Thr500 | 1X Tyr41(P)<br>3X Tyr41(H)<br>1X Asn330(H)<br>2X Asp355(H)<br>2X Arg357(H) | | Missing<br>1X Tyr41(H)<br><br>1X Asp355(H)<br>1X Arg357(H)<br>1X Leu45(H) | | | | | | | | |
| Asn501 | 3X Tyr41(H)<br>1X Lys353(H) | | | 2X Tyr41(H)<br>3X Lys353(H) | 1X Tyr41(H)<br>3X Lys353(H) | 5X Tyr41(H)<br>6X Lys353(H) | | | | | |
| Gly502 | 1X Lys353(P)<br>1X Lys353(H)<br>2X Gly354(H) | | | | | | 2X Lys353(H)<br>4X Gly354(H)<br>1X Thr324(H) | 2X Lys353(H)<br>4X Gly354(H) | 4X Gly354(H)<br>3X Thr324(H) | | |
| Tyr505 | 5X Lys353(H)<br>1X Gly354(H) | | | | | | | | | 2X Lys353(H)<br>Missing | 7X Lys353(H)<br>Missing<br>_**1X Lys353 (*)**_ |
