## Supplementary Table 1 for "A computational approach to evaluate the combined effect of SARS-CoV-2 RBD mutations and ACE2 receptor genetic variants on infectivity: The COVID-19 host-pathogen nexus"

**Supplementary Table 2**: Changes in the polar and hydrophobic interactions in 24 different SARS-CoV-2 spike RBD mutations that occurred in the 18 contact residues with nine hACE2 genetic variants. P: polar, H: hydrophobic. Only missing, changed and new interactions are mentioned for each variant. (*)= salt bridge.

| Spike RBD Wuhan strain contact residues | hACE2 isoform 1 contact residues | Wuhan Strain | | | | | | | |
|---|---|---|---|---|---|---|---|---|---|
| | | D355A | D355N | E35D | E35K | F40L | M82I | T27A | S19P |
| Lys417 | 1X Asp30(P)<br>1X Asp30(H) | | | | | | | | |
| Gly446 | 1X Gln42(P) | | | | | | | | |
| Tyr449 | 1X Asp38(P)<br>1X Gln42(P)<br>3X Asp38(H) | | | | | | | | |
| Tyr453 | 1X His34(P)<br>2X His34(H) | | | | | | | | |
| Leu455 | 4X His34(H) | | | | | | | | |
| Phe456 | 1X Thr27(H)<br>1X Asp30(H) | | | | | | | 1X Ala27(H) | |
| Ala475 | 1X Ser19(P)<br>2X Ser19(H)<br>1X Gln24(H) | | | | | | | | Missing<br>1XPro19(H) |
| Gly476 | 1X Ser19(H) | | | | | | | | 1XPro19(H) |
| Phe486 | 1X Met82(H)<br>4X Tyr83(H) | | | | | | 1X Ile82(H) | | |
| Asn487 | 1X Gln24(P)<br>1X Tyr83(P)<br>6X Gln24(H)<br>3X Tyr83(H) | | | | | | | | |
| Tyr489 | 1X Thr27(H)<br>1X Phe28(H) | | | | | | | Messing | |
| Gln493 | 2X His34(H)<br>1X Glu35(H) | | | 1XAsp35(P)<br>1X Asp35(H) | 1X Lys35(H) | | | | |
| Gly496 | 1X Lys353(P)<br>1X Asp38(H)<br>2X Lys353(H) | | | | | | | | |
| Gln498 | 1X Gln42(P)<br>3X Tyr41(H)<br>2X Gln42(H)<br>1X Leu45(H) | | | | | | | | |
| Thr500 | X Tyr41(P)<br>3X Tyr41(H)<br>1X Asn330(H)<br>2X Asp355(H)<br>2X Arg357(H) | 1X Ala355(H) | 1X Asn355(P)<br>4x Asn355(H) | | | | | | |
| Asn501 | 3X Tyr41(H)<br>1X Lys353(H) | | 1X Asn355(H) | | | | | | |
| Gly502 | 1X Lys353(P)<br>1X Lys353(H)<br>2X Gly354(H) | | | | | | | | |
| Tyr505 | 5X Lys353(H)<br>1X Gly354(H) | | | | | | | | |

**Supplementary Table 2**: (Continued)

| Spike RBD Wuhan strain contact residues | hACE2 isoform 1 contact residues | K417N | | | | | | | |
|---|---|---|---|---|---|---|---|---|---|
| | | D355A | D355N | E35D | E35K | F40L | M82I | T27A | S19P |
| Lys417 | 1X Asp30(P)<br>1X Asp30(H) | Missing<br>Missing | Missing<br>Missing | Missing<br>Missing | Missing<br>Missing | Missing<br>Missing | Missing<br>Missing | Missing<br>Missing | Missing<br>Missing |
| Gly446 | 1X Gln42(P) | | | | | | | | |
| Tyr449 | 1X Asp38(P)<br>1X Gln42(P)<br>3X Asp38(H) | | | | | | | | |
| Tyr453 | 1X His34(P)<br>2X His34(H) | | | | | | | | |
| Leu455 | 4X His34(H) | | | | | | | | |
| Phe456 | 1X Thr27(H)<br>1X Asp30(H) | | | | | | | | |
| Ala475 | 1X Ser19(P)<br>2X Ser19(H)<br>1X Gln24(H) | | | | | | | | Missing<br>1XPro19(H) |
| Gly476 | 1X Ser19(H) | | | | | | | | |
| Phe486 | 1X Met82(H)<br>4X Tyr83(H) | | | | | | 1X Ile82(H) | | |
| Asn487 | 1X Gln24(P)<br>1X Tyr83(P)<br>6X Gln24(H)<br>3X Tyr83(H) | | | | | | | | |
| Tyr489 | 1X Thr27(H)<br>1X Phe28(H) | | | | | | | Missing | |
| Gln493 | 2X His34(H)<br>1X Glu35(H) | | | 1X Asp35(P)<br>1X Asp35(H) | 1X Lys35(H) | | | | |
| Gly496 | 1X Lys353(P)<br>1X Asp38(H)<br>2X Lys353(H) | | | | | | | | |
| Gln498 | 1X Gln42(P)<br>3X Tyr41(H)<br>2X Gln42(H)<br>1X Leu45(H) | | | | | | | | |
| Thr500 | 1X Tyr41(P)<br>3X Tyr41(H)<br>1X Asn330(H)<br>2X Asp355(H)<br>2X Arg357(H) | 1X Ala355(H) | 1X Asn355(P)<br>4X Ala355(H) | | | | | | |
| Asn501 | 3X Tyr41(H)<br>1X Lys353(H) | | 1X Ala355(H) | | | | | | |
| Gly502 | 1X Lys353(P)<br>1X Lys353(H)<br>2X Gly354(H) | | | | | | | | |
| Tyr505 | 5X Lys353(H)<br>1X Gly354(H) | | | | | | | | |

**Supplementary Table 2**: (Continued)

| Spike RBD Wuhan strain contact residues | hACE2 isoform 1 contact residues | K417R | | | | | | | |
|---|---|---|---|---|---|---|---|---|---|
| | | D355A | D355N | E35D | E35K | F40L | M82I | T27A | S19P |
| Lys417 | 1X Asp30(P)<br>1X Asp30(H) | 4X Asp30(H) | 4X Asp30(H) | 4X Asp30(H) | 4X Asp30(H) | 4X Asp30(H) | 4X Asp30(H) | 4X Asp30(H) | 4X Asp30(H) |
| Gly446 | 1X Gln42(P) | | | | | | | | |
| Tyr449 | 1X Asp38(P)<br>1X Gln42(P)<br>3X Asp38(H) | | | | | | | | |
| Tyr453 | 1X His34(P)<br>2X His34(H) | | | | | | | | |
| Leu455 | 4X His34(H) | | | | | | | | |
| Phe456 | 1X Thr27(H)<br>1X Asp30(H) | | | | | | | 1X Ala7(H) | |
| Ala475 | 1X Ser19(P)<br>2X Ser19(H)<br>1X Gln24(H) | | | | | | | | Missing<br>1X Pro19(H) |
| Gly476 | 1X Ser19(H) | | | | | | | | 1X Pro19(H) |
| Phe486 | 1X Met82(H)<br>4X Tyr83(H) | | | | | | 1X Ile82(H) | | |
| Asn487 | 1X Gln24(P)<br>1X Tyr83(P)<br>6X Gln24(H)<br>3X Tyr83(H) | | | | | | | | |
| Tyr489 | 1X Thr27(H)<br>1X Phe28(H) | | | | | | | Missing | |
| Gln493 | 2X His34(H)<br>1X Glu35(H) | | | 1X Asp35(P)<br>1X Asp35(H) | 1X Lys35(H) | | | | |
| Gly496 | 1X Lys353(P)<br>1X Asp38(H)<br>2X Lys353(H) | | | | | | | | |
| Gln498 | 1X Gln42(P)<br>3X Tyr41(H)<br>2X Gln42(H)<br>1X Leu45(H) | | | | | | | | |
| Thr500 | 1X Tyr41(P)<br>3X Tyr41(H)<br>1X Asn330(H)<br>2X Asp355(H)<br>2X Arg357(H) | 1X Ala355(H) | 1X Asn355(P)<br>4X Asn355(H) | | | | | | |
| Asn501 | 3X Tyr41(H)<br>1X Lys353(H) | | 1X Asn355(H) | | | | | | |
| Gly502 | 1X Lys353(P)<br>1X Lys353(H)<br>2X Gly354(H) | | | | | | | | |
| Tyr505 | 5X Lys353(H)<br>1X Gly354(H) | | | | | | | | |

**Supplementary Table 2**: (Continued)

| Spike RBD Wuhan strain contact residues | hACE2 isoform 1 contact residues | G446A | | | | | | | |
|---|---|---|---|---|---|---|---|---|---|
| | | D355A | D355N | E35D | E35K | F40L | M82I | T27A | S19P |
| Lys417 | 1X Asp30(P)<br>1X Asp30(H) | | | | | | | | |
| Gly446 | 1X Gln42(P) | Missing | Missing | Missing | Missing | Missing | Missing | Missing | Missing |
| Tyr449 | 1X Asp38(P)<br>1X Gln42(P)<br>3X Asp38(H) | | | | | | | | |
| Tyr453 | 1X His34(P)<br>2X His34(H) | | | | | | | | |
| Leu455 | 4X His34(H) | | | | | | | | |
| Phe456 | 1X Thr27(H)<br>1X Asp30(H) | | | | | | | 1X Ala27(H) | |
| Ala475 | 1X Ser19(P)<br>2X Ser19(H)<br>1X Gln24(H) | | | | | | | | Missing<br>1X Pro19(H) |
| Gly476 | 1X Ser19(H) | | | | | | | | 1X Pro19(H) |
| Phe486 | 1X Met82(H)<br>4X Tyr83(H) | | | | | | 1X Ile82(H) | | |
| Asn487 | 1X Gln24(P)<br>1X Tyr83(P)<br>6X Gln24(H)<br>3X Tyr83(H) | | | | | | | | |
| Tyr489 | 1X Thr27(H)<br>1X Phe28(H) | | | | | | | Missing | |
| Gln493 | 2X His34(H)<br>1X Glu35(H) | | | 1X Asp35 (P)<br>1X Asp35(H) | 1X Lys35(H) | | | | |
| Gly496 | 1X Lys353(P)<br>1X Asp38(H)<br>2X Lys353(H) | | | | | | | | |
| Gln498 | 1X Gln42(P)<br>3X Tyr41(H)<br>2X Gln42(H)<br>1X Leu45(H) | | | | | | | | |
| Thr500 | 1X Tyr41(P)<br>3X Tyr41(H)<br>1X Asn330(H)<br>2X Asp355(H)<br>2X Arg357(H) | 1X Aal355(H) | 1X Asn355(P)<br>4X Aal355(H) | | | | | | |
| Asn501 | 3X Tyr41(H)<br>1X Lys353(H) | | 1X Aal355(H) | | | | | | |
| Gly502 | 1X Lys353(P)<br>1X Lys353(H)<br>2X Gly354(H) | | | | | | | | |
| Tyr505 | 5X Lys353(H)<br>1X Gly354(H) | | | | | | | | |

**Supplementary Table 2**: (Continued)

| Spike RBD Wuhan strain contact residues | hACE2 isoform 1 contact residues | Y449N | | | | | | | |
|---|---|---|---|---|---|---|---|---|---|
| | | D355A | D355N | E35D | E35K | F40L | M82I | T27A | S19P |
| Lys417 | 1X Asp30(P)<br>1X Asp30(H) | | | | | | | | |
| Gly446 | 1X Gln42(P) | | | | | | | | |
| Tyr449 | 1X Asp38(P)<br>1X Gln42(P)<br>3X Asp38(H) | Missing<br>Missing<br>Missing | Missing<br>Missing<br>Missing | Missing<br>Missing<br>Missing | Missing<br>Missing<br>Missing | Missing<br>Missing<br>Missing | Missing<br>Missing<br>Missing | Missing<br>Missing<br>Missing | Missing<br>Missing<br>Missing |
| Tyr453 | 1X His34(P)<br>2X His34(H) | | | | | | | | |
| Leu455 | 4X His34(H) | | | | | | | | |
| Phe456 | 1X Thr27(H)<br>1X Asp30(H) | | | | | | | 1X Ala27(H) | |
| Ala475 | 1X Ser19(P)<br>2X Ser19(H)<br>1X Gln24(H) | | | | | | | | Missing<br>1X Pro19(H) |
| Gly476 | 1X Ser19(H) | | | | | | | | 1X Pro19(H) |
| Phe486 | 1X Met82(H)<br>4X Tyr83(H) | | | | | | 1X Ile82(H) | | |
| Asn487 | 1X Gln24(P)<br>1X Tyr83(P)<br>6X Gln24(H)<br>3X Tyr83(H) | | | | | | | | |
| Tyr489 | 1X Thr27(H)<br>1X Phe28(H) | | | | | | | Missing | |
| Gln493 | 2X His34(H)<br>1X Glu35(H) | | | 1X Asp35 (P)<br>1X Asp35(H) | 1X Lys35(H) | | | | |
| Gly496 | 1X Lys353(P)<br>1X Asp38(H)<br>2X Lys353(H) | | | | | | | | |
| Gln498 | 1X Gln42(P)<br>3X Tyr41(H)<br>2X Gln42(H)<br>1X Leu45(H) | | | | | | | | |
| Thr500 | 1X Tyr41(P)<br>3X Tyr41(H)<br>1X Asn330(H)<br>2X Asp355(H)<br>2X Arg357(H) | 1X Ala355(H) | 1X Asn355 (P)<br>4X Asn355(H) | | | | | | |
| Asn501 | 3X Tyr41(H)<br>1X Lys353(H) | | 1X Asn355(H) | | | | | | |
| Gly502 | 1X Lys353(P)<br>1X Lys353(H)<br>2X Gly354(H) | | | | | | | | |
| Tyr505 | 5X Lys353(H)<br>1X Gly354(H) | | | | | | | | |

**Supplementary Table 2**: (Continued)

| Spike RBD Wuhan strain contact residues | hACE2 isoform 1 contact residues | Y453F | | | | | | | |
|---|---|---|---|---|---|---|---|---|---|
| | | D355A | D355N | E35D | E35K | F40L | M82I | T27A | S19P |
| Lys417 | 1X Asp30(P)<br>1X Asp30(H) | | | | | | | | |
| Gly446 | 1X Gln42(P) | | | | | | | | |
| Tyr449 | 1X Asp38(P)<br>1X Gln42(P)<br>3X Asp38(H) | | | | | | | | |
| Tyr453 | 1X His34(P)<br>2X His34(H) | Missing<br>Missing | Missing<br>Missing | Missing<br>Missing | Missing<br>Missing | Missing<br>Missing | Missing<br>Missing | Missing<br>Missing | Missing<br>Missing |
| Leu455 | 4X His34(H) | 1X His34(H) | 1X His34(H) | 1X His34(H) | 1X His34(H) | 1X His34(H) | 1X His34(H) | 1X His34(H) | 1X His34(H) |
| Phe456 | 1X Thr27(H)<br>1X Asp30(H) | | | | | | | 1X Ala27(H) | |
| Ala475 | 1X Ser19(P)<br>2X Ser19(H)<br>1X Gln24(H) | | | | | | | | Missing<br>1X Pro19(H) |
| Gly476 | 1X Ser19(H) | | | | | | | | 1X Pro19(H) |
| Phe486 | 1X Met82(H)<br>4X Tyr83(H) | | | | | | 1X Ile82(H) | | |
| Asn487 | 1X Gln24(P)<br>1X Tyr83(P)<br>6X Gln24(H)<br>3X Tyr83(H) | | | | | | | | |
| Tyr489 | 1X Thr27(H)<br>1X Phe28(H) | | | | | | | Missing | |
| Gln493 | 2X His34(H)<br>1X Glu35(H) | 1X His34(H) | 1X His34(H) | 1X Asp35(P)<br>1X His34(H)<br>1X Asp35(H) | 1X His34(H)<br>1X lys35(H) | 1X His34(H) | 1X His34(H) | 1X His34(H) | 1X His34(H) |
| Gly496 | 1X Lys353(P)<br>1X Asp38(H)<br>2X Lys353(H) | | | | | | | | |
| Gln498 | 1X Gln42(P)<br>3X Tyr41(H)<br>2X Gln42(H)<br>1X Leu45(H) | | | | | | | | |
| Thr500 | 1X Tyr41(P)<br>3X Tyr41(H)<br>1X Asn330(H)<br>2X Asp355(H)<br>2X Arg357(H) | 1X Aal355(H) | 1X Asn355(P)<br>4X Aal355(H) | | | | | | |
| Asn501 | 3X Tyr41(H)<br>1X Lys353(H) | | 1X Aal355(H) | | | | | | |
| Gly502 | 1X Lys353(P)<br>1X Lys353(H)<br>2X Gly354(H) | | | | | | | | |
| Tyr505 | 5X Lys353(H)<br>1X Gly354(H) | | | | | | | | |

**Supplementary Table 2**: (Continued)

| Spike RBD Wuhan strain contact residues | hACE2 isoform 1 contact residues | L455F | | | | | | | |
|---|---|---|---|---|---|---|---|---|---|
| | | D355A | D355N | E35D | E35K | F40L | M82I | T27A | S19P |
| Lys417 | 1X Asp30(P)<br>1X Asp30(H) | | | | | | | | |
| Gly446 | 1X Gln42(P) | | | | | | | | |
| Tyr449 | 1X Asp38(P)<br>1X Gln42(P)<br>3X Asp38(H) | | | | | | | | |
| Tyr453 | 1X His34(P)<br>2X His34(H) | | | | | | | | |
| Leu455 | 4X His34(H) | 4X Asp30(H)<br>1X Lys 31(H) | 4X Asp30(H)<br>1X Lys 31(H) | 4X Asp30(H)<br>1X Lys 31(H) | 4X Asp30(H)<br>1X Lys 31(H) | 4X Asp30(H)<br>1X Lys 31(H) | 4X Asp30(H)<br>1X Lys 31(H) | 4X Asp30(H)<br>1X Lys 31(H) | 4X Asp30(H)<br>1X Lys 31(H) |
| Phe456 | 1X Thr27(H)<br>1X Asp30(H) | | | | | | | 1X Ala27(H) | |
| Ala475 | 1X Ser19(P)<br>2X Ser19(H)<br>1X Gln24(H) | | | | | | | | Missing<br>1X Pro19(H) |
| Gly476 | 1X Ser19(H) | | | | | | | | 1X Pro19(H) |
| Phe486 | 1X Met82(H)<br>4X Tyr83(H) | | | | | | 1X Ile82(H) | | |
| Asn487 | 1X Gln24(P)<br>1X Tyr83(P)<br>6X Gln24(H)<br>3X Tyr83(H) | | | | | | | | |
| Tyr489 | 1X Thr27(H)<br>1X Phe28(H) | | | | | | | Missing | |
| Gln493 | 2X His34(H)<br>1X Glu35(H) | | | 1X Asp35(P)<br>1X Asp35(H) | 1X Lys35(H) | | | | |
| Gly496 | 1X Lys353(P)<br>1X Asp38(H)<br>2X Lys353(H) | | | | | | | | |
| Gln498 | 1X Gln42(P)<br>3X Tyr41(H)<br>2X Gln42(H)<br>1X Leu45(H) | | | | | | | | |
| Thr500 | 1X Tyr41(P)<br>3X Tyr41(H)<br>1X Asn330(H)<br>2X Asp355(H)<br>2X Arg357(H) | 1X Ala355(H) | 1X Asn335(P)<br>4X Asn355(H) | | | | | | |
| Asn501 | 3X Tyr41(H)<br>1X Lys353(H) | | 1X Asn355(H) | | | | | | |
| Gly502 | 1X Lys353(P)<br>1X Lys353(H)<br>2X Gly354(H) | | | | | | | | |
| Tyr505 | 5X Lys353(H)<br>1X Gly354(H) | | | | | | | | |

**Supplementary Table 2**: (Continued)

| Spike RBD Wuhan strain contact residues | hACE2 isoform 1 contact residues | F456L | | | | | | | |
|---|---|---|---|---|---|---|---|---|---|
| | | D355A | D355N | E35D | E35K | F40L | M82I | T27A | S19P |
| Lys417 | 1X Asp30(P)<br>1X Asp30(H) | | | | | | | | |
| Gly446 | 1X Gln42(P) | | | | | | | | |
| Tyr449 | 1X Asp38(P)<br>1X Gln42(P)<br>3X Asp38(H) | | | | | | | | |
| Tyr453 | 1X His34(P)<br>2X His34(H) | | | | | | | | |
| Leu455 | 4X His34(H) | | | | | | | | |
| Phe456 | 1X Thr27(H)<br>1X Asp30(H) | Missing | Missing | Missing | Missing | Missing | Missing | Missing<br>Missing | Missing |
| Ala475 | 1X Ser19(P)<br>2X Ser19(H)<br>1X Gln24(H) | | | | | | | | Missing<br>1X Pro19(H) |
| Gly476 | 1X Ser19(H) | | | | | | | | 1X Pro19(H) |
| Phe486 | 1X Met82(H)<br>4X Tyr83(H) | | | | | | 1X Ile82(H) | | |
| Asn487 | 1X Gln24(P)<br>1X Tyr83(P)<br>6X Gln24(H)<br>3X Tyr83(H) | | | | | | | | |
| Tyr489 | 1X Thr27(H)<br>1X Phe28(H) | | | | | | | Missing | |
| Gln493 | 2X His34(H)<br>1X Glu35(H) | | | 1X Asp35(P)<br>1X Asp35(H) | 1X Lys35(H) | | | | |
| Gly496 | 1X Lys353(P)<br>1X Asp38(H)<br>2X Lys353(H) | | | | | | | | |
| Gln498 | 1X Gln42(P)<br>3X Tyr41(H)<br>2X Gln42(H)<br>1X Leu45(H) | | | | | | | | |
| Thr500 | 1X Tyr41(P)<br>3X Tyr41(H)<br>1X Asn330(H)<br>2X Asp355(H)<br>2X Arg357(H) | 1X Ala355(H) | 1X Asn355(P)<br>4X Asn355(H) | | | | | | |
| Asn501 | 3X Tyr41(H)<br>1X Lys353(H) | | 1X Ala355(H) | | | | | | |
| Gly502 | 1X Lys353(P)<br>1X Lys353(H)<br>2X Gly354(H) | | | | | | | | |
| Tyr505 | 5X Lys353(H)<br>1X Gly354(H) | | | | | | | | |

**Supplementary Table 2**: (Continued)

| Spike RBD Wuhan strain contact residues | hACE2 isoform 1 contact residues | A475V | | | | | | | |
|---|---|---|---|---|---|---|---|---|---|
| | | D355A | D355N | E35D | E35K | F40L | M82I | T27A | S19P |
| Lys417 | 1X Asp30(P)<br>1X Asp30(H) | | | | | | | | |
| Gly446 | 1X Gln42(P) | | | | | | | | |
| Tyr449 | 1X Asp38(P)<br>1X Gln42(P)<br>3X Asp38(H) | | | | | | | | |
| Tyr453 | 1X His34(P)<br>2X His34(H) | | | | | | | | |
| Leu455 | 4X His34(H) | | | | | | | | |
| Phe456 | 1X Thr27(H)<br>1X Asp30(H) | | | | | | | | |
| Ala475 | 1X Ser19(P)<br>2X Ser19(H)<br>1X Gln24(H) | 2X Gln24(H)<br>2X Thr27(H) | 2X Gln24(H)<br>2X Thr27(H) | 2X Gln24(H)<br>2X Thr27(H) | 2X Gln24(H)<br>2X Thr27(H) | 2X Gln24(H)<br>2X Thr27(H) | 2X Gln24(H)<br>2X Thr27(H) | 2X Gln24(H)<br>1X Thr27(H) | 1X Pro19(H)<br>2X Gln24(H)<br>1X Thr27(H) |
| Gly476 | 1X Ser19(H) | | | | | | | | 1X Pro19(H) |
| Phe486 | 1X Met82(H)<br>4X Tyr83(H) | | | | | | 1X Ile82(H) | | |
| Asn487 | 1X Gln24(P)<br>1X Tyr83(P)<br>6X Gln24(H)<br>3X Tyr83(H) | | | | | | | | |
| Tyr489 | 1X Thr27(H)<br>1X Phe28(H) | | | | | | | | |
| Gln493 | 2X His34(H)<br>1X Glu35(H) | | | 1X Asp35(P)<br>1X Asp35(H) | 1X Lys35(H) | | | Missing | |
| Gly496 | 1X Lys353(P)<br>1X Asp38(H)<br>2X Lys353(H) | | | | | | | | |
| Gln498 | 1X Gln42(P)<br>3X Tyr41(H)<br>2X Gln42(H)<br>1X Leu45(H) | | | | | | | | |
| Thr500 | 1X Tyr41(P)<br>3X Tyr41(H)<br>1X Asn330(H)<br>2X Asp355(H)<br>2X Arg357(H) | 1X Ala355(H) | 1X Asn355(P)<br>4X Asn355(H) | | | | | | |
| Asn501 | 3X Tyr41(H)<br>1X Lys353(H) | | 1X Asn355(H) | | | | | | |
| Gly502 | 1X Lys353(P)<br>1X Lys353(H)<br>2X Gly354(H) | | | | | | | | |
| Tyr505 | 5X Lys353(H)<br>1X Gly354(H) | | | | | | | | |

**Supplementary Table 2**: (Continued)

| Spike RBD Wuhan strain contact residues | hACE2 isoform 1 contact residues | G476S | | | | | | | |
|---|---|---|---|---|---|---|---|---|---|
| | | **D355A** | **D355N** | **E35D** | **E35K** | **F40L** | **M82I** | **T27A** | **S19P** |
| Lys417 | 1X Asp30(P)<br>1X Asp30(H) | | | | | | | | |
| Gly446 | 1X Gln42(P) | | | | | | | | |
| Tyr449 | 1X Asp38(P)<br>1X Gln42(P)<br>3X Asp38(H) | | | | | | | | |
| Tyr453 | 1X His34(P)<br>2X His34(H) | | | | | | | | |
| Leu455 | 4X His34(H) | | | | | | | | |
| Phe456 | 1X Thr27(H)<br>1X Asp30(H) | | | | | | | 1X Ala27(H) | |
| Ala475 | 1X Ser19(P)<br>2X Ser19(H)<br>1X Gln24(H) | | | | | | | | Missing<br>1X Pro19(H) |
| Gly476 | 1X Ser19(H) | Missing | Missing | Missing | Missing | Missing | Missing | Missing | Missing |
| Phe486 | 1X Met82(H)<br>4X Tyr83(H) | | | | | | 1X Ile82(H) | | |
| Asn487 | 1X Gln24(P)<br>1X Tyr83(P)<br>6X Gln24(H)<br>3X Tyr83(H) | | | | | | | | |
| Tyr489 | 1X Thr27(H)<br>1X Phe28(H) | | | | | | | Missing | |
| Gln493 | 2X His34(H)<br>1X Glu35(H) | | | 1X Asp35(P)<br>1X Asp35(H) | 1X Lys35(H) | | | | |
| Gly496 | 1X Lys353(P)<br>1X Asp38(H)<br>2X Lys353(H) | | | | | | | | |
| Gln498 | 1X Gln42(P)<br>3X Tyr41(H)<br>2X Gln42(H)<br>1X Leu45(H) | | | | | | | | |
| Thr500 | 1X Tyr41(P)<br>3X Tyr41(H)<br>1X Asn330(H)<br>2X Asp355(H)<br>2X Arg357(H) | 1X Ala355(H) | 1X Asn335(P)<br>4X Asn355(H) | | | | | | |
| Asn501 | 3X Tyr41(H)<br>1X Lys353(H) | | 1X Asn355(H) | | | | | | |
| Gly502 | 1X Lys353(P)<br>1X Lys353(H)<br>2X Gly354(H) | | | | | | | | |
| Tyr505 | 5X Lys353(H)<br>1X Gly354(H) | | | | | | | | |

**Supplementary Table 2**: (Continued)

| Spike RBD Wuhan strain contact residues | hACE2 isoform 1 contact residues | G476A | | | | | | | |
|---|---|---|---|---|---|---|---|---|---|
| | | D355A | D355N | E35D | E35K | F40L | M82I | T27A | S19P |
| Lys417 | 1X Asp30(P)<br>1X Asp30(H) | | | | | | | | |
| Gly446 | 1X Gln42(P) | | | | | | | | |
| Tyr449 | 1X Asp38(P)<br>1X Gln42(P)<br>3X Asp38(H) | | | | | | | | |
| Tyr453 | 1X His34(P)<br>2X His34(H) | | | | | | | | |
| Leu455 | 4X His34(H) | | | | | | | | |
| Phe456 | 1X Thr27(H)<br>1X Asp30(H) | | | | | | | 1X Ala27(H) | |
| Ala475 | 1X Ser19(P)<br>2X Ser19(H)<br>1X Gln24(H) | | | | | | | | Missing<br>1X Pro19(H) |
| Gly476 | 1X Ser19(H) | Missing | Missing | Missing | Missing | Missing | Missing | Missing | Missing |
| Phe486 | 1X Met82(H)<br>4X Tyr83(H) | | | | | | 1X Ile82(H) | | |
| Asn487 | 1X Gln24(P)<br>1X Tyr83(P)<br>6X Gln24(H)<br>3X Tyr83(H) | | | | | | | | |
| Tyr489 | 1X Thr27(H)<br>1X Phe28(H) | | | | | | | Missing | |
| Gln493 | 2X His34(H)<br>1X Glu35(H) | | | 1X Asp35(P)<br>1X Asp35(H) | 1X Lys35(H) | | | | |
| Gly496 | 1X Lys353(P)<br>1X Asp38(H)<br>2X Lys353(H) | | | | | | | | |
| Gln498 | 1X Gln42(P)<br>3X Tyr41(H)<br>2X Gln42(H)<br>1X Leu45(H) | | | | | | | | |
| Thr500 | 1X Tyr41(P)<br>3X Tyr41(H)<br>1X Asn330(H)<br>2X Asp355(H)<br>2X Arg357(H) | 1X Ala355(H) | 1X Asn335(P)<br>4X Asn355(H) | | | | | | |
| Asn501 | 3X Tyr41(H)<br>1X Lys353(H) | | 1X Asn355(H) | | | | | | |
| Gly502 | 1X Lys353(P)<br>1X Lys353(H)<br>2X Gly354(H) | | | | | | | | |
| Tyr505 | 5X Lys353(H)<br>1X Gly354(H) | | | | | | | | |

**Supplementary Table 2**: (Continued)

| Spike RBD Wuhan strain contact residues | hACE2 isoform 1 contact residues | F486L | | | | | | | |
|---|---|---|---|---|---|---|---|---|---|
| | | D355A | D355N | E35D | E35K | F40L | M82I | T27A | S19P |
| Lys417 | 1X Asp30(P)<br>1X Asp30(H) | | | | | | | | |
| Gly446 | 1X Gln42(P) | | | | | | | | |
| Tyr449 | 1X Asp38(P)<br>1X Gln42(P)<br>3X Asp38(H) | | | | | | | | |
| Tyr453 | 1X His34(P)<br>2X His34(H) | | | | | | | | |
| Leu455 | 4X His34(H) | | | | | | | | |
| Phe456 | 1X Thr27(H)<br>1X Asp30(H) | | | | | | | 1X Ala27(H) | |
| Ala475 | 1X Ser19(P)<br>2X Ser19(H)<br>1X Gln24(H) | | | | | | | | Missing<br>1X Pro19(H) |
| Gly476 | 1X Ser19(H) | | | | | | | | 1X Pro19(H) |
| Phe486 | 1X Met82(H)<br>4X Tyr83(H) | 1X Tyr83(H) | 1X Tyr83(H) | 1X Tyr83(H) | 1X Tyr83(H) | 1X Tyr83(H) | 1X Ile82(H)<br>1X Tyr83(H) | 1X Tyr83(H) | 1X Tyr83(H) |
| Asn487 | 1X Gln24(P)<br>1X Tyr83(P)<br>6X Gln24(H)<br>3X Tyr83(H) | | | | | | | | |
| Tyr489 | 1X Thr27(H)<br>1X Phe28(H) | | | | | | | Missing | |
| Gln493 | 2X His34(H)<br>1X Glu35(H) | | | 1X Asp35(P)<br>1X Asp35(H) | 1X Lys35(H) | | | | |
| Gly496 | 1X Lys353(P)<br>1X Asp38(H)<br>2X Lys353(H) | | | | | | | | |
| Gln498 | 1X Gln42(P)<br>3X Tyr41(H)<br>2X Gln42(H)<br>1X Leu45(H) | | | | | | | | |
| Thr500 | 1X Tyr41(P)<br>3X Tyr41(H)<br>1X Asn330(H)<br>2X Asp355(H)<br>2X Arg357(H) | 1X Asp355(H) | 1X Asn355(P)<br>4X Asn355(H) | | | | | | |
| Asn501 | 3X Tyr41(H)<br>1X Lys353(H) | | 1X Asn355(H) | | | | | | |
| Gly502 | 1X Lys353(P)<br>1X Lys353(H)<br>2X Gly354(H) | | | | | | | | |
| Tyr505 | 5X Lys353(H)<br>1X Gly354(H) | | | | | | | | |

**Supplementary Table 2**: (Continued)

| Spike RBD Wuhan strain contact residues | hACE2 isoform 1 contact residues | Q493R | | | | | | | |
|---|---|---|---|---|---|---|---|---|---|
| | | **D355A** | **D355N** | **E35D** | **E35K** | **F40L** | **M82I** | **T27A** | **S19P** |
| Lys417 | 1X Asp30(P)<br>1X Asp30(H) | | | | | | | | |
| Gly446 | 1X Gln42(P) | | | | | | | | |
| Tyr449 | 1X Asp38(P)<br>1X Gln42(P)<br>3X Asp38(H) | | | | | | | | |
| Tyr453 | 1X His34(P)<br>2X His34(H) | | | | | | | | |
| Leu455 | 4X His34(H) | | | | | | | | |
| Phe456 | 1X Thr27(H)<br>1X Asp30(H) | | | | | | | 1X Ala27(H) | |
| Ala475 | 1X Ser19(P)<br>2X Ser19(H)<br>1X Gln24(H) | | | | | | | | Missing<br>1X Pr019(H) |
| Gly476 | 1X Ser19(H) | | | | | | | | 1X Pro19(H) |
| Phe486 | 1X Met82(H)<br>4X Tyr83(H) | | | | | | 1X Ile82(H) | | |
| Asn487 | 1X Gln24(P)<br>1X Tyr83(P)<br>6X Gln24(H)<br>3X Tyr83(H) | | | | | | | | |
| Tyr489 | 1X Thr27(H)<br>1X Phe28(H) | | | | | | | Missing | |
| Gln493 | 2X His34(H)<br>1X Glu35(H) | 1X Glu35 (P)<br>Missing<br>4X Glu35(H) | 1X Glu35 (P)<br>Missing<br>4X Glu35(H) | Missing<br>Missing<br>***1X Asp35 (*)*** | Missing<br>Missing | 1X Glu35 (P)<br>Missing<br>4X Glu35(H) | 1X Glu35 (P)<br>Missing<br>4X Glu35(H) | 1X Glu35 (P)<br>Missing<br>4X Glu35(H) | 1X Glu35 (P)<br>Missing<br>4X Glu35(H) |
| Gly496 | 1X Lys353(P)<br>1X Asp38(H)<br>2X Lys353(H) | | | | | | | | |
| Gln498 | 1X Gln42(P)<br>3X Tyr41(H)<br>2X Gln42(H)<br>1X Leu45(H) | | | | | | | | |
| Thr500 | 1X Tyr41(P)<br>3X Tyr41(H)<br>1X Asn330(H)<br>2X Asp355(H)<br>2X Arg357(H) | 1X Asp355(H) | 1X Asn355(P)<br>4X Asn355(H) | | | | | | |
| Asn501 | 3X Tyr41(H)<br>1X Lys353(H) | | 1X Asn355(H) | | | | | | |
| Gly502 | 1X Lys353(P)<br>1X Lys353(H)<br>2X Gly354(H) | | | | | | | | |
| Tyr505 | 5X Lys353(H)<br>1X Gly354(H) | | | | | | | | |

**Supplementary Table 2**: (Continued)

| Spike RBD Wuhan strain contact residues | hACE2 isoform 1 contact residues | Q493L | | | | | | | |
|---|---|---|---|---|---|---|---|---|---|
| | | D355A | D355N | E35D | E35K | F40L | M82I | T27A | S19P |
| Lys417 | 1X Asp30(P)<br>1X Asp30(H) | | | | | | | | |
| Gly446 | 1X Gln42(P) | | | | | | | | |
| Tyr449 | 1X Asp38(P)<br>1X Gln42(P)<br>3X Asp38(H) | | | | | | | | |
| Tyr453 | 1X His34(P)<br>2X His34(H) | | | | | | | | |
| Leu455 | 4X His34(H) | | | | | | | | |
| Phe456 | 1X Thr27(H)<br>1X Asp30(H) | | | | | | | 1X Ala27(H) | |
| Ala475 | 1X Ser19(P)<br>2X Ser19(H)<br>1X Gln24(H) | | | | | | | | Missing<br>1X Pr019(H) |
| Gly476 | 1X Ser19(H) | | | | | | | | 1X Pro19(H) |
| Phe486 | 1X Met82(H)<br>4X Tyr83(H) | | | | | | 1X Ile82(H) | | |
| Asn487 | 1X Gln24(P)<br>1X Tyr83(P)<br>6X Gln24(H)<br>3X Tyr83(H) | | | | | | | | |
| Tyr489 | 1X Thr27(H)<br>1X Phe28(H) | | | | | | | Missing | |
| Gln493 | 2X His34(H)<br>1X Glu35(H) | 1X His34(H)<br>Missing<br>1X Lys31(H) | 1X His34(H)<br>Missing<br>1X Lys31(H) | 1X His34(H)<br>Missing<br>1X lys31(H) | 1X His34(H)<br>Missing<br>1X lys31(H) | 1X His34(H)<br>Missing<br>1X Lys31(H) | 1X His34(H)<br>Missing<br>1X Lys31(H) | 1X His34(H)<br>Missing<br>1X Lys31(H) | 1X His34(H)<br>Missing<br>1X Lys31(H) |
| Gly496 | 1X Lys353(P)<br>1X Asp38(H)<br>2X Lys353(H) | | | | | | | | |
| Gln498 | 1X Gln42(P)<br>3X Tyr41(H)<br>2X Gln42(H)<br>1X Leu45(H) | | | | | | | | |
| Thr500 | 1X Tyr41(P)<br>3X Tyr41(H)<br>1X Asn330(H)<br>2X Asp355(H)<br>2X Arg357(H) | 1X Asp355(H) | 1X Asn355(P)<br>4X Asn355(H) | | | | | | |
| Asn501 | 3X Tyr41(H)<br>1X Lys353(H) | | 1X Asn355(H) | | | | | | |
| Gly502 | 1X Lys353(P)<br>1X Lys353(H)<br>2X Gly354(H) | | | | | | | | |
| Tyr505 | 5X Lys353(H)<br>1X Gly354(H) | | | | | | | | |

**Supplementary Table 2**: (Continued)

| Spike RBD Wuhan strain contact residues | hACE2 isoform 1 contact residues | G496C | | | | | | | |
|---|---|---|---|---|---|---|---|---|---|
| | | D355A | D355N | E35D | E35K | F40L | M82I | T27A | S19P |
| Lys417 | 1X Asp30(P)<br>1X Asp30(H) | | | | | | | | |
| Gly446 | 1X Gln42(P) | | | | | | | | |
| Tyr449 | 1X Asp38(P)<br>1X Gln42(P)<br>3X Asp38(H) | | | | | | | | |
| Tyr453 | 1X His34(P)<br>2X His34(H) | | | | | | | | |
| Leu455 | 4X His34(H) | | | | | | | | |
| Phe456 | 1X Thr27(H)<br>1X Asp30(H) | | | | | | | 1X Ala27(H) | |
| Ala475 | 1X Ser19(P)<br>2X Ser19(H)<br>1X Gln24(H) | | | | | | | | Missing<br>1X Pr019(H) |
| Gly476 | 1X Ser19(H) | | | | | | | | 1X Pro19(H) |
| Phe486 | 1X Met82(H)<br>4X Tyr83(H) | | | | | | 1X Ile82(H) | | |
| Asn487 | 1X Gln24(P)<br>1X Tyr83(P)<br>6X Gln24(H)<br>3X Tyr83(H) | | | | | | | | |
| Tyr489 | 1X Thr27(H)<br>1X Phe28(H) | | | | | | | Missing | |
| Gln493 | 2X His34(H)<br>1X Glu35(H) | | | 1XAsp35(P)<br>1X Asp35(H) | 1X Lys35(H) | | | | |
| Gly496 | 1X Lys353(P)<br>1X Asp38(H)<br>2X Lys353(H) | 7X Asp38(H) | 7X Asn38(H) | 7X Asp38(H) | 7X Asp38(H) | 7X Asp38(H) | 7X Asp38(H) | 7X Asp38(H) | 7X Asp38(H) |
| Gln498 | 1X Gln42(P)<br>3X Tyr41(H)<br>2X Gln42(H)<br>1X Leu45(H) | | | | | | | | |
| Thr500 | 1X Tyr41(P)<br>3X Tyr41(H)<br>1X Asn330(H)<br>2X Asp355(H)<br>2X Arg357(H) | 1X Asp355(H) | 1X Asn355(P)<br>4X Asn355(H) | | | | | | |
| Asn501 | 3X Tyr41(H)<br>1X Lys353(H) | | 1X Asn355(H) | | | | | | |
| Gly502 | 1X Lys353(P)<br>1X Lys353(H)<br>2X Gly354(H) | | | | | | | | |
| Tyr505 | 5X Lys353(H)<br>1X Gly354(H) | | | | | | | | |

**Supplementary Table 2**: (Continued)

| Spike RBD Wuhan strain contact residues | hACE2 isoform 1 contact residues | T500I | | | | | | | |
|---|---|---|---|---|---|---|---|---|---|
| | | D355A | D355N | E35D | E35K | F40L | M82I | T27A | S19P |
| Lys417 | 1X Asp30(P)<br>1X Asp30(H) | | | | | | | | |
| Gly446 | 1X Gln42(P) | | | | | | | | |
| Tyr449 | 1X Asp38(P)<br>1X Gln42(P)<br>3X Asp38(H) | | | | | | | | |
| Tyr453 | 1X His34(P)<br>2X His34(H) | | | | | | | | |
| Leu455 | 4X His34(H) | | | | | | | | |
| Phe456 | 1X Thr27(H)<br>1X Asp30(H) | | | | | | | 1X Ala27(H) | |
| Ala475 | 1X Ser19(P)<br>2X Ser19(H)<br>1X Gln24(H) | | | | | | | | Missing<br>1X Pr019(H) |
| Gly476 | 1X Ser19(H) | | | | | | | | 1X Pro19(H) |
| Phe486 | 1X Met82(H)<br>4X Tyr83(H) | | | | | | 1X Ile82(H) | | |
| Asn487 | 1X Gln24(P)<br>1X Tyr83(P)<br>6X Gln24(H)<br>3X Tyr83(H) | | | | | | | | |
| Tyr489 | 1X Thr27(H)<br>1X Phe28(H) | | | | | | | Missing | |
| Gln493 | 2X His34(H)<br>1X Glu35(H) | | | 1XAsp35(P)<br>1X Asp35(H) | 1X Lys35(H) | | | | |
| Gly496 | 1X Lys353(P)<br>1X Asp38(H)<br>2X Lys353(H) | | | | | | | | |
| Gln498 | 1X Gln42(P)<br>3X Tyr41(H)<br>2X Gln42(H)<br>1X Leu45(H) | 2X Tyr41(H) | 2X Tyr41(H) | 2X Tyr41(H) | 2X Tyr41(H) | 2X Tyr41(H) | 2X Tyr41(H) | 2X Tyr41(H) | 2X Tyr41(H) |
| Thr500 | 1X Tyr41(P)<br>3X Tyr41(H)<br>1X Asn330(H)<br>2X Asp355(H)<br>2X Arg357(H) | Missing<br>1X Tyr41(H)<br>Missing<br>1X Arg357(H)<br>1X Leu45(H) | Missing<br>1X Tyr41(H)<br>1X Asp355(H)<br>1X Arg357(H)<br>1X Leu45(H) | Missing<br>1X Tyr41(H)<br>1X Asp355(H)<br>1X Arg357(H)<br>1X Leu45(H) | Missing<br>1X Tyr41(H)<br>1X Asp355(H)<br>1X Arg357(H)<br>1X Leu45(H) | Missing<br>1X Tyr41(H)<br>1X Asp355(H)<br>1X Arg357(H)<br>1X Leu45(H) | Missing<br>1X Tyr41(H)<br>1X Asp355(H)<br>1X Arg357(H)<br>1X Leu45(H) | Missing<br>1X Tyr41(H)<br>1X Asp355(H)<br>1X Arg357(H)<br>1X Leu45(H) | Missing<br>1X Tyr41(H)<br>1X Asp355(H)<br>1X Arg357(H)<br>1X Leu45(H) |
| Asn501 | 3X Tyr41(H)<br>1X Lys353(H) | 1X Tyr41(H) | 1X Tyr41(H)<br>1X Asn355(H) | 1X Tyr41(H) | 1X Tyr41(H) | 1X Tyr41(H) | 1X Tyr41(H) | 1X Tyr41(H) | 1X Tyr41(H) |
| Gly502 | 1X Lys353(P)<br>1X Lys353(H)<br>2X Gly354(H) | | | | | | | | |
| Tyr505 | 5X Lys353(H)<br>1X Gly354(H) | | | | | | | | |

**Supplementary Table 2**: (Continued)

| Spike RBD Wuhan strain contact residues | hACE2 isoform 1 contact residues | N501T | | | | | | | |
|---|---|---|---|---|---|---|---|---|---|
| | | D355A | D355N | E35D | E35K | F40L | M82I | T27A | S19P |
| Lys417 | 1X Asp30(P)<br>1X Asp30(H) | | | | | | | | |
| Gly446 | 1X Gln42(P) | | | | | | | | |
| Tyr449 | 1X Asp38(P)<br>1X Gln42(P)<br>3X Asp38(H) | | | | | | | | |
| Tyr453 | 1X His34(P)<br>2X His34(H) | | | | | | | | |
| Leu455 | 4X His34(H) | | | | | | | | |
| Phe456 | 1X Thr27(H)<br>1X Asp30(H) | | | | | | | 1X Ala27(H) | |
| Ala475 | 1X Ser19(P)<br>2X Ser19(H)<br>1X Gln24(H) | | | | | | | | Missing<br>1X Pr019(H) |
| Gly476 | 1X Ser19(H) | | | | | | | | 1X Pro19(H) |
| Phe486 | 1X Met82(H)<br>4X Tyr83(H) | | | | | | 1X Ile82(H) | | |
| Asn487 | 1X Gln24(P)<br>1X Tyr83(P)<br>6X Gln24(H)<br>3X Tyr83(H) | | | | | | | | |
| Tyr489 | 1X Thr27(H)<br>1X Phe28(H) | | | | | | | Missing | |
| Gln493 | 2X His34(H)<br>1X Glu35(H) | | | 1XAsp35(P)<br>1X Asp35(H) | 1X Lys35(H) | | | | |
| Gly496 | 1X Lys353(P)<br>1X Asp38(H)<br>2X Lys353(H) | | | | | | | | |
| Gln498 | 1X Gln42(P)<br>3X Tyr41(H)<br>2X Gln42(H)<br>1X Leu45(H) | | | | | | | | |
| Thr500 | 1X Tyr41(P)<br>3X Tyr41(H)<br>1X Asn330(H)<br>2X Asp355(H)<br>2X Arg357(H) | 1X Ala355(H) | 1X Asn355(P)<br>4X Asn355(H) | | | | | | |
| Asn501 | 3X Tyr41(H)<br>1X Lys353(H) | 2X Tyr41(H)<br>3X Lys353(H) | 2X Tyr41(H)<br>3X Lys353(H)<br>1XAsn355(H) | 2X Tyr41(H)<br>3X Lys353(H) | 2X Tyr41(H)<br>3X Lys353(H) | 2X Tyr41(H)<br>3X Lys353(H) | 2X Tyr41(H)<br>3X Lys353(H) | 2X Tyr41(H)<br>3X Lys353(H) | 2X Tyr41(H)<br>3X Lys353(H) |
| Gly502 | 1X Lys353(P)<br>1X Lys353(H)<br>2X Gly354(H) | | | | | | | | |
| Tyr505 | 5X Lys353(H)<br>1X Gly354(H) | | | | | | | | |

**Supplementary Table 2**: (Continued)

| Spike RBD Wuhan strain contact residues | hACE2 isoform 1 contact residues | N501S | | | | | | | |
|---|---|---|---|---|---|---|---|---|---|
| | | D355A | D355N | E35D | E35K | F40L | M82I | T27A | S19P |
| Lys417 | 1X Asp30(P)<br>1X Asp30(H) | | | | | | | | |
| Gly446 | 1X Gln42(P) | | | | | | | | |
| Tyr449 | 1X Asp38(P)<br>1X Gln42(P)<br>3X Asp38(H) | | | | | | | | |
| Tyr453 | 1X His34(P)<br>2X His34(H) | | | | | | | | |
| Leu455 | 4X His34(H) | | | | | | | | |
| Phe456 | 1X Thr27(H)<br>1X Asp30(H) | | | | | | | 1X Ala27(H) | |
| Ala475 | 1X Ser19(P)<br>2X Ser19(H)<br>1X Gln24(H) | | | | | | | | Missing<br>1X Pr019(H) |
| Gly476 | 1X Ser19(H) | | | | | | | | 1X Pro19(H) |
| Phe486 | 1X Met82(H)<br>4X Tyr83(H) | | | | | | 1X Ile82(H) | | |
| Asn487 | 1X Gln24(P)<br>1X Tyr83(P)<br>6X Gln24(H)<br>3X Tyr83(H) | | | | | | | | |
| Tyr489 | 1X Thr27(H)<br>1X Phe28(H) | | | | | | | Missing | |
| Gln493 | 2X His34(H)<br>1X Glu35(H) | | | 1XAsp35(P)<br>1X Asp35(H) | 1X Lys35(H) | | | | |
| Gly496 | 1X Lys353(P)<br>1X Asp38(H)<br>2X Lys353(H) | | | | | | | | |
| Gln498 | 1X Gln42(P)<br>3X Tyr41(H)<br>2X Gln42(H)<br>1X Leu45(H) | | | | | | | | |
| Thr500 | 1X Tyr41(P)<br>3X Tyr41(H)<br>1X Asn330(H)<br>2X Asp355(H)<br>2X Arg357(H) | 1X Ala355(H) | 1X Asn355(P)<br>4X Asn355(H) | | | | | | |
| Asn501 | 3X Tyr41(H)<br>1X Lys353(H) | 1X Tyr41(H)<br>3X Lys353(H) | 1X Tyr41(H)<br>3X Lys353(H)<br>1XAsn355(H) | 1X Tyr41(H)<br>3X Lys353(H) | 1X Tyr41(H)<br>3X Lys353(H) | 1X Tyr41(H)<br>3X Lys353(H) | 1X Tyr41(H)<br>3X Lys353(H) | 1X Tyr41(H)<br>3X Lys353(H) | 1X Tyr41(H)<br>3X Lys353(H) |
| Gly502 | 1X Lys353(P)<br>1X Lys353(H)<br>2X Gly354(H) | | | | | | | | |
| Tyr505 | 5X Lys353(H)<br>1X Gly354(H) | | | | | | | | |

**Supplementary Table 2**: (Continued)

| Spike RBD Wuhan strain contact residues | hACE2 isoform 1 contact residues | N501Y | | | | | | | |
|---|---|---|---|---|---|---|---|---|---|
| | | D355A | D355N | E35D | E35K | F40L | M82I | T27A | S19P |
| Lys417 | 1X Asp30(P)<br>1X Asp30(H) | | | | | | | | |
| Gly446 | 1X Gln42(P) | | | | | | | | |
| Tyr449 | 1X Asp38(P)<br>1X Gln42(P)<br>3X Asp38(H) | | | | | | | | |
| Tyr453 | 1X His34(P)<br>2X His34(H) | | | | | | | | |
| Leu455 | 4X His34(H) | | | | | | | | |
| Phe456 | 1X Thr27(H)<br>1X Asp30(H) | | | | | | | 1X Ala27(H) | |
| Ala475 | 1X Ser19(P)<br>2X Ser19(H)<br>1X Gln24(H) | | | | | | | | Missing<br>1X Pr019(H) |
| Gly476 | 1X Ser19(H) | | | | | | | | 1X Pro19(H) |
| Phe486 | 1X Met82(H)<br>4X Tyr83(H) | | | | | | 1X Ile82(H) | | |
| Asn487 | 1X Gln24(P)<br>1X Tyr83(P)<br>6X Gln24(H)<br>3X Tyr83(H) | | | | | | | | |
| Tyr489 | 1X Thr27(H)<br>1X Phe28(H) | | | | | | | Missing | |
| Gln493 | 2X His34(H)<br>1X Glu35(H) | | | 1XAsp35(P)<br>1X Asp35(H) | 1X Lys35(H) | | | | |
| Gly496 | 1X Lys353(P)<br>1X Asp38(H)<br>2X Lys353(H) | | | | | | | | |
| Gln498 | 1X Gln42(P)<br>3X Tyr41(H)<br>2X Gln42(H)<br>1X Leu45(H) | | | | | | | | |
| Thr500 | 1X Tyr41(P)<br>3X Tyr41(H)<br>1X Asn330(H)<br>2X Asp355(H)<br>2X Arg357(H) | 1X Ala355(H) | 1X Asn355(P)<br>4X Asn355(H) | | | | | | |
| Asn501 | 3X Tyr41(H)<br>1X Lys353(H) | 5X Tyr41(H)<br>6X Lys353(H) | 5X Tyr41(H)<br>6X Lys353(H)<br>1XAsn355(H) | 5X Tyr41(H)<br>6X Lys353(H) | 5X Tyr41(H)<br>6X Lys353(H) | 5X Tyr41(H)<br>6X Lys353(H) | 5X Tyr41(H)<br>6X Lys353(H) | 5X Tyr41(H)<br>6X Lys353(H) | 5X Tyr41(H)<br>6X Lys353(H) |
| Gly502 | 1X Lys353(P)<br>1X Lys353(H)<br>2X Gly354(H) | | | | | | | | |
| Tyr505 | 5X Lys353(H)<br>1X Gly354(H) | | | | | | | | |

**Supplementary Table 2**: (Continued)

| Spike RBD Wuhan strain contact residues | hACE2 isoform 1 contact residues | G502D | | | | | | | |
|---|---|---|---|---|---|---|---|---|---|
| | | D355A | D355N | E35D | E35K | F40L | M82I | T27A | S19P |
| Lys417 | 1X Asp30(P)<br>1X Asp30(H) | | | | | | | | |
| Gly446 | 1X Gln42(P) | | | | | | | | |
| Tyr449 | 1X Asp38(P)<br>1X Gln42(P)<br>3X Asp38(H) | | | | | | | | |
| Tyr453 | 1X His34(P)<br>2X His34(H) | | | | | | | | |
| Leu455 | 4X His34(H) | | | | | | | | |
| Phe456 | 1X Thr27(H)<br>1X Asp30(H) | | | | | | | 1X Ala27(H) | |
| Ala475 | 1X Ser19(P)<br>2X Ser19(H)<br>1X Gln24(H) | | | | | | | | Missing<br>1X Pr019(H) |
| Gly476 | 1X Ser19(H) | | | | | | | | 1X Pro19(H) |
| Phe486 | 1X Met82(H)<br>4X Tyr83(H) | | | | | | 1X Ile82(H) | | |
| Asn487 | 1X Gln24(P)<br>1X Tyr83(P)<br>6X Gln24(H)<br>3X Tyr83(H) | | | | | | | | |
| Tyr489 | 1X Thr27(H)<br>1X Phe28(H) | | | | | | | Missing | |
| Gln493 | 2X His34(H)<br>1X Glu35(H) | | | 1XAsp35(P)<br>1X Asp35(H) | 1X Lys35(H) | | | | |
| Gly496 | 1X Lys353(P)<br>1X Asp38(H)<br>2X Lys353(H) | | | | | | | | |
| Gln498 | 1X Gln42(P)<br>3X Tyr41(H)<br>2X Gln42(H)<br>1X Leu45(H) | | | | | | | | |
| Thr500 | 1X Tyr41(P)<br>3X Tyr41(H)<br>1X Asn330(H)<br>2X Asp355(H)<br>2X Arg357(H) | 1X Ala355(H) | 1X Asn355(P)<br>4X Asn355(H) | | | | | | |
| Asn501 | 3X Tyr41(H)<br>1X Lys353(H) | | 1XAsn355(H) | | | | | | |
| Gly502 | 1X Lys353(P)<br>1X Lys353(H)<br>2X Gly354(H) | 2X Lys353(H)<br>4X Gly354(H)<br>1X Thr324(H) | 2X Lys353(H)<br>4X Gly354(H)<br>1X Thr324(H) | 2X Lys353(H)<br>4X Gly354(H)<br>1X Thr324(H) | 2X Lys353(H)<br>4X Gly354(H)<br>1X Thr324(H) | 2X Lys353(H)<br>4X Gly354(H)<br>1X Thr324(H) | 2X Lys353(H)<br>4X Gly354(H)<br>1X Thr324(H) | 2X Lys353(H)<br>4X Gly354(H)<br>1X Thr324(H) | 2X Lys353(H)<br>4X Gly354(H)<br>1X Thr324(H) |
| Tyr505 | 5X Lys353(H)<br>1X Gly354(H) | | | | | | | | |

**Supplementary Table 2**: (Continued)

| Spike RBD Wuhan strain contact residues | hACE2 isoform 1 contact residues | G502C | | | | | | | |
|---|---|---|---|---|---|---|---|---|---|
| | | D355A | D355N | E35D | E35K | F40L | M82I | T27A | S19P |
| Lys417 | 1X Asp30(P)<br>1X Asp30(H) | | | | | | | | |
| Gly446 | 1X Gln42(P) | | | | | | | | |
| Tyr449 | 1X Asp38(P)<br>1X Gln42(P)<br>3X Asp38(H) | | | | | | | | |
| Tyr453 | 1X His34(P)<br>2X His34(H) | | | | | | | | |
| Leu455 | 4X His34(H) | | | | | | | | |
| Phe456 | 1X Thr27(H)<br>1X Asp30(H) | | | | | | | 1X Ala27(H) | |
| Ala475 | 1X Ser19(P)<br>2X Ser19(H)<br>1X Gln24(H) | | | | | | | | Missing<br>1X Pr019(H) |
| Gly476 | 1X Ser19(H) | | | | | | | | 1X Pro19(H) |
| Phe486 | 1X Met82(H)<br>4X Tyr83(H) | | | | | | 1X Ile82(H) | | |
| Asn487 | 1X Gln24(P)<br>1X Tyr83(P)<br>6X Gln24(H)<br>3X Tyr83(H) | | | | | | | | |
| Tyr489 | 1X Thr27(H)<br>1X Phe28(H) | | | | | | | Missing | |
| Gln493 | 2X His34(H)<br>1X Glu35(H) | | | 1XAsp35(P)<br>1X Asp35(H) | 1X Lys35(H) | | | | |
| Gly496 | 1X Lys353(P)<br>1X Asp38(H)<br>2X Lys353(H) | | | | | | | | |
| Gln498 | 1X Gln42(P)<br>3X Tyr41(H)<br>2X Gln42(H)<br>1X Leu45(H) | | | | | | | | |
| Thr500 | 1X Tyr41(P)<br>3X Tyr41(H)<br>1X Asn330(H)<br>2X Asp355(H)<br>2X Arg357(H) | 1X Ala355(H) | 1X Asn355(P)<br>4X Asn355(H) | | | | | | |
| Asn501 | 3X Tyr41(H)<br>1X Lys353(H) | | 1XAsn355(H) | | | | | | |
| Gly502 | 1X Lys353(P)<br>1X Lys353(H)<br>2X Gly354(H) | 2X Lys353(H)<br>4X Gly354(H) | 2X Lys353(H)<br>4X Gly354(H) | 2X Lys353(H)<br>4X Gly354(H) | 2X Lys353(H)<br>4X Gly354(H) | 2X Lys353(H)<br>4X Gly354(H) | 2X Lys353(H)<br>4X Gly354(H) | 2X Lys353(H)<br>4X Gly354(H) | 2X Lys353(H)<br>4X Gly354(H) |
| Tyr505 | 5X Lys353(H)<br>1X Gly354(H) | | | | | | | | |

**Supplementary Table 2**: (Continued)

| Spike RBD Wuhan strain contact residues | hACE2 isoform 1 contact residues | G502R | | | | | | | |
|---|---|---|---|---|---|---|---|---|---|
| | | D355A | D355N | E35D | E35K | F40L | M82I | T27A | S19P |
| Lys417 | 1X Asp30(P)<br>1X Asp30(H) | | | | | | | | |
| Gly446 | 1X Gln42(P) | | | | | | | | |
| Tyr449 | 1X Asp38(P)<br>1X Gln42(P)<br>3X Asp38(H) | | | | | | | | |
| Tyr453 | 1X His34(P)<br>2X His34(H) | | | | | | | | |
| Leu455 | 4X His34(H) | | | | | | | | |
| Phe456 | 1X Thr27(H)<br>1X Asp30(H) | | | | | | | 1X Ala27(H) | |
| Ala475 | 1X Ser19(P)<br>2X Ser19(H)<br>1X Gln24(H) | | | | | | | | Missing<br>1X Pr019(H) |
| Gly476 | 1X Ser19(H) | | | | | | | | 1X Pro19(H) |
| Phe486 | 1X Met82(H)<br>4X Tyr83(H) | | | | | | 1X Ile82(H) | | |
| Asn487 | 1X Gln24(P)<br>1X Tyr83(P)<br>6X Gln24(H)<br>3X Tyr83(H) | | | | | | | | |
| Tyr489 | 1X Thr27(H)<br>1X Phe28(H) | | | | | | | Missing | |
| Gln493 | 2X His34(H)<br>1X Glu35(H) | | | 1XAsp35(P)<br>1X Asp35(H) | 1X Lys35(H) | | | | |
| Gly496 | 1X Lys353(P)<br>1X Asp38(H)<br>2X Lys353(H) | | | | | | | | |
| Gln498 | 1X Gln42(P)<br>3X Tyr41(H)<br>2X Gln42(H)<br>1X Leu45(H) | | | | | | | | |
| Thr500 | 1X Tyr41(P)<br>3X Tyr41(H)<br>1X Asn330(H)<br>2X Asp355(H)<br>2X Arg357(H) | 1X Ala355(H) | 1X Asn355(P)<br>4X Asn355(H) | | | | | | |
| Asn501 | 3X Tyr41(H)<br>1X Lys353(H) | | 1XAsn355(H) | | | | | | |
| Gly502 | 1X Lys353(P)<br>1X Lys353(H)<br>2X Gly354(H) | 4X Gly354(H)<br>3X Thr324(H) | 4X Gly354(H)<br>3X Thr324(H) | 4X Gly354(H)<br>3X Thr324(H) | 4X Gly354(H)<br>3X Thr324(H) | 4X Gly354(H)<br>3X Thr324(H) | 4X Gly354(H)<br>3X Thr324(H) | 4X Gly354(H)<br>3X Thr324(H) | 4X Gly354(H)<br>3X Thr324(H) |
| Tyr505 | 5X Lys353(H)<br>1X Gly354(H) | | | | | | | | |

**Supplementary Table 2**: (Continued)

| Spike RBD Wuhan strain contact residues | hACE2 isoform 1 contact residues | Y505H | | | | | | | |
|---|---|---|---|---|---|---|---|---|---|
| | | D355A | D355N | E35D | E35K | F40L | M82I | T27A | S19P |
| Lys417 | 1X Asp30(P)<br>1X Asp30(H) | | | | | | | | |
| Gly446 | 1X Gln42(P) | | | | | | | | |
| Tyr449 | 1X Asp38(P)<br>1X Gln42(P)<br>3X Asp38(H) | | | | | | | | |
| Tyr453 | 1X His34(P)<br>2X His34(H) | | | | | | | | |
| Leu455 | 4X His34(H) | | | | | | | | |
| Phe456 | 1X Thr27(H)<br>1X Asp30(H) | | | | | | | 1X Ala27(H) | |
| Ala475 | 1X Ser19(P)<br>2X Ser19(H)<br>1X Gln24(H) | | | | | | | | Missing<br>1X Pr019(H) |
| Gly476 | 1X Ser19(H) | | | | | | | | 1X Pro19(H) |
| Phe486 | 1X Met82(H)<br>4X Tyr83(H) | | | | | | 1X Ile82(H) | | |
| Asn487 | 1X Gln24(P)<br>1X Tyr83(P)<br>6X Gln24(H)<br>3X Tyr83(H) | | | | | | | | |
| Tyr489 | 1X Thr27(H)<br>1X Phe28(H) | | | | | | | Missing | |
| Gln493 | 2X His34(H)<br>1X Glu35(H) | | | 1XAsp35(P)<br>1X Asp35(H) | 1X Lys35(H) | | | | |
| Gly496 | 1X Lys353(P)<br>1X Asp38(H)<br>2X Lys353(H) | | | | | | | | |
| Gln498 | 1X Gln42(P)<br>3X Tyr41(H)<br>2X Gln42(H)<br>1X Leu45(H) | | | | | | | | |
| Thr500 | 1X Tyr41(P)<br>3X Tyr41(H)<br>1X Asn330(H)<br>2X Asp355(H)<br>2X Arg357(H) | 1X Asp355(H) | 1X Asn355(P)<br>4X Asn355(H) | | | | | | |
| Asn501 | 3X Tyr41(H)<br>1X Lys353(H) | | 1X Asn355(H) | | | | | | |
| Gly502 | 1X Lys353(P)<br>1X Lys353(H)<br>2X Gly354(H) | | | | | | | | |
| Tyr505 | 5X Lys353(H)<br>1X Gly354(H) | 2X Lys353(H)<br>Missing | 2X Lys353(H)<br>Missing | 2X Lys353(H)<br>Missing | 2X Lys353(H)<br>Missing | 2X Lys353(H)<br>Missing | 2X Lys353(H)<br>Missing | 2X Lys353(H)<br>Missing | 2X Lys353(H)<br>Missing |

**Supplementary Table 2**: (Continued)

| Spike RBD Wuhan strain contact residues | hACE2 isoform 1 contact residues | Y505E | | | | | | | |
|---|---|---|---|---|---|---|---|---|---|
| | | D355A | D355N | E35D | E35K | F40L | M82I | T27A | S19P |
| Lys417 | 1X Asp30(P)<br>1X Asp30(H) | | | | | | | | |
| Gly446 | 1X Gln42(P) | | | | | | | | |
| Tyr449 | 1X Asp38(P)<br>1X Gln42(P)<br>3X Asp38(H) | | | | | | | | |
| Tyr453 | 1X His34(P)<br>2X His34(H) | | | | | | | | |
| Leu455 | 4X His34(H) | | | | | | | | |
| Phe456 | 1X Thr27(H)<br>1X Asp30(H) | | | | | | | 1X Ala27(H) | |
| Ala475 | 1X Ser19(P)<br>2X Ser19(H)<br>1X Gln24(H) | | | | | | | | Missing<br>1X Pr019(H) |
| Gly476 | 1X Ser19(H) | | | | | | | | 1X Pro19(H) |
| Phe486 | 1X Met82(H)<br>4X Tyr83(H) | | | | | | 1X Ile82(H) | | |
| Asn487 | 1X Gln24(P)<br>1X Tyr83(P)<br>6X Gln24(H)<br>3X Tyr83(H) | | | | | | | | |
| Tyr489 | 1X Thr27(H)<br>1X Phe28(H) | | | | | | | Missing | |
| Gln493 | 2X His34(H)<br>1X Glu35(H) | | | 1XAsp35(P)<br>1X Asp35(H) | 1X Lys35(H) | | | | |
| Gly496 | 1X Lys353(P)<br>1X Asp38(H)<br>2X Lys353(H) | | | | | | | | |
| Gln498 | 1X Gln42(P)<br>3X Tyr41(H)<br>2X Gln42(H)<br>1X Leu45(H) | | | | | | | | |
| Thr500 | 1X Tyr41(P)<br>3X Tyr41(H)<br>1X Asn330(H)<br>2X Asp355(H)<br>2X Arg357(H) | 1X Asp355(H) | 1X Asn355(P)<br>4X Asn355(H) | | | | | | |
| Asn501 | 3X Tyr41(H)<br>1X Lys353(H) | | 1X Asn355(H) | | | | | | |
| Gly502 | 1X Lys353(P)<br>1X Lys353(H)<br>2X Gly354(H) | | | | | | | | |
| Tyr505 | 5X Lys353(H)<br>1X Gly354(H) | 7X Lys353(H)<br>Missing<br>***1X Lys353 (*)*** | 7X Lys353(H)<br>Missing<br>***1X Lys353 (*)*** | 7X Lys353(H)<br>Missing<br>***1X Lys353 (*)*** | 7X Lys353(H)<br>Missing<br>***1X Lys353 (*)*** | 7X Lys353(H)<br>Missing<br>***1X Lys353 (*)*** | 7X Lys353(H)<br>Missing<br>***1X Lys353 (*)*** | 7X Lys353(H)<br>Missing<br>***1X Lys353 (*)*** | 7X Lys353(H)<br>Missing<br>***1X Lys353 (*)*** |
